# Acetylation-Primed SUMOylation Drives RORβ Turnover via a p300-SIRT1 Regulatory Axis

**DOI:** 10.1101/2024.10.30.621067

**Authors:** Timothy R. O’Leary, Denis Shutin, Nadeska I. Montalvan, Nereida M. Abad-Yang, Vuong Dang, Dean P. Edwards, Patrick R. Griffin, Mi Ra Chang

## Abstract

Retinoic acid receptor-related orphan receptor beta (RORβ) is a transcription factor expressed in the central nervous system, retina, and bone that regulates circadian rhythms, retinal neurogenesis, and inflammatory signaling. Despite these critical functions, the mechanisms governing RORβ stability remain poorly understood. Here, we identify a post-translational regulatory axis in which the lysine acetyltransferase p300 and the NAD⁺-dependent deacetylase SIRT1 control RORβ stability and transcriptional activity. p300-mediated acetylation increases RORβ abundance, while SIRT1 modulates turnover through both catalytic and non-catalytic scaffolding mechanisms. K176 acetylation in the hinge primes UBC9/PIAS1-mediated SUMOylation at nearby K179, marking RORβ for proteasomal degradation and reducing transcriptional output, providing a mechanistic framework for targeting RORβ in neurological and retinal disorders, and bone homeostasis.

**Highlights:**

- p300 acetylates RORβ at eight lysines; SIRT1 reverses this via catalytic activity
- K176 acetylation primes UBC9/PIAS1-mediated SUMOylation at nearby K179
- SUMOylated RORβ undergoes proteasomal degradation leading to reduced RORβ-mediated transcriptional output
- p300 displaces ubiquitin E3 ligases from RORβ; SIRT1 activation reduces PIAS1 association

**In Brief:** O’Leary et al. define a post-translational circuit in which p300 acetylates K176 within the RORβ hinge domain, priming SUMOylation at nearby K179 preferentially by UBC9/PIAS1. SUMOylated RORβ is targeted for proteasomal degradation with diminished transcriptional activity, while SIRT1 deacetylase activity antagonizes this pathway via both catalytic activity and protein-protein interactions to stabilize active RORβ.

Graphical Abstract

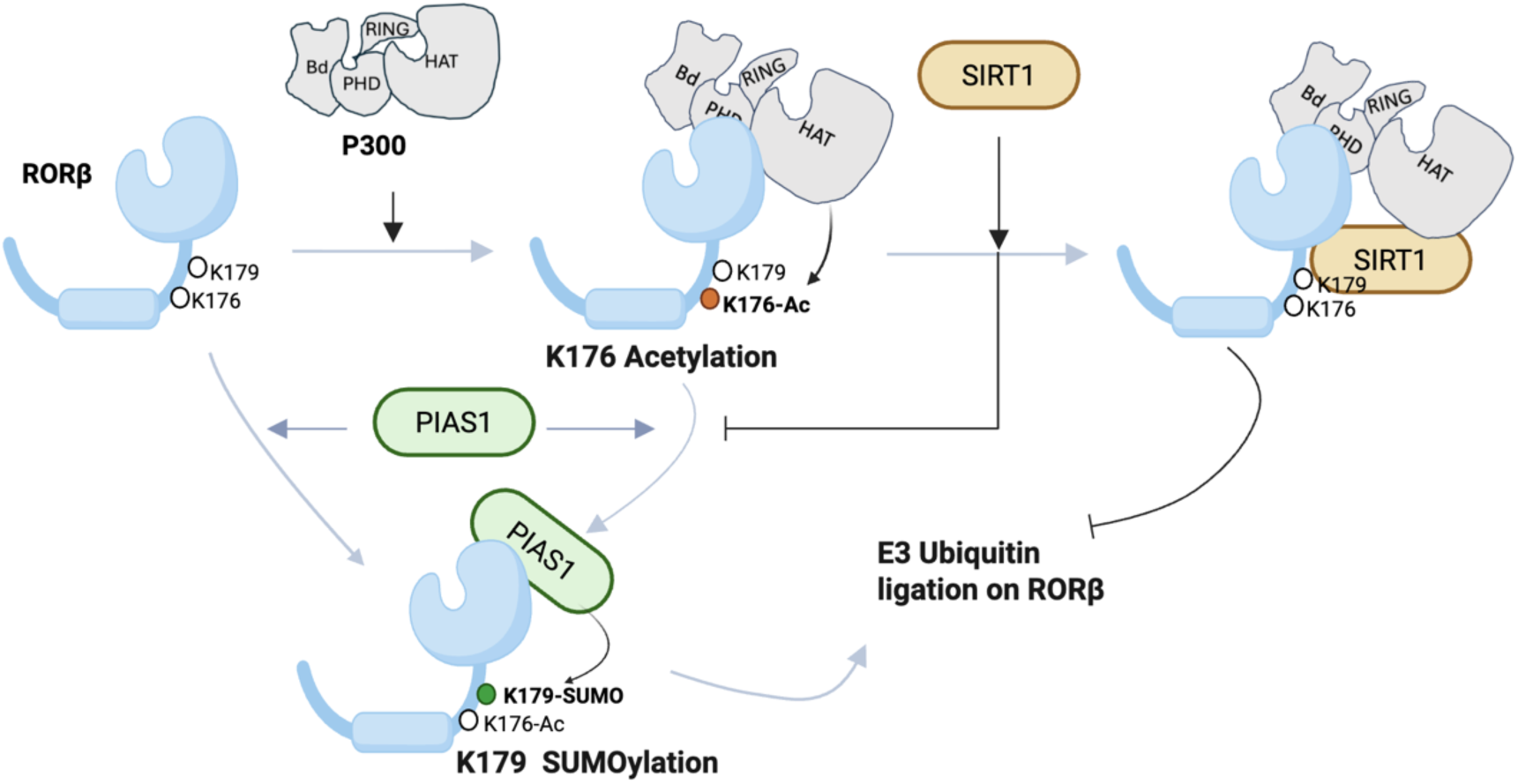

## Introduction

The nuclear receptor (NR) superfamily comprises ligand-regulated transcription factors that govern essential aspects of mammalian physiology and represent a major class of therapeutic targets. Among these, the retinoic acid receptor-related orphan receptors (RORs), members of the NR1F subfamily, regulate diverse processes including glucose and lipid metabolism, circadian rhythms, bone homeostasis, immune function, retinal development, and neural circuit function ^1–6^. The ROR subfamily consists of three genes, RORα, RORβ, and RORγ, sharing structural homology with retinoic acid receptors (RARs) ^7^. While RORα and RORγ are modulated by oxysterols and cholesterol metabolites ^8–10^, RORβ remains enigmatic: although it binds oxysterols and can be antagonized by all-trans retinoic acid (ATRA), its endogenous ligand and direct transcriptional targets remain undefined ^11,12^. RORβ has been significantly less studied as the field focused on RORα and RORγ ligands modulating immune function as potential therapeutics for autoimmune disease and cancer ^7,13^.

RORβ is expressed as two major isoforms, RORβ1 and RORβ2, generated by alternative promoter usage differing only in their N-terminus, with RORβ2 primarily expressed in the retina and RORβ1 more broadly distributed throughout the central nervous system, including cortical and sensory regions ^14,15 16–18^. In the CNS, RORβ regulates sensory processing and circadian rhythms; its deletion disrupts rhythmicity and retinal development resulting in blindness ^16,17,19–22 23^. Dysregulation of RORβ has been linked to sleep disorders, bipolar disorder, schizophrenia, epilepsy, and Alzheimer’s disease ^24,25^. In bone, RORβ suppresses osteoblast differentiation and mineralization, implicating it in osteoarthritis and age-related bone loss ^26^. Emerging evidence suggests RORβ functions as a tumor suppressor in neuroblastoma, colorectal, gastric, and head and neck cancers, where low expression correlates with poor clinical outcomes ^27–30^. Despite these critical roles, RORβ remains understudied relative to RORα and RORγ, with key gaps in our understanding of its PTMs, endogenous ligands, transcriptional programs, and context-dependent functions.

RORβ adopts the canonical domain architecture of the NR superfamily, comprising a variable, intrinsically disordered N-terminal A/B domain that contains the ligand-independent activation function-1 (AF-1), followed by a conserved two-zinc finger DNA-binding domain (DBD). The DBD is connected via a flexible hinge region to the C-terminal ligand-binding domain (LBD), which harbors the ligand-dependent AF-2 coregulator interface ^31^. The hinge region of NRs is often enriched in post-translational modifications (PTMs), including acetylation, SUMOylation, ubiquitination, and phosphorylation, which are known to influence receptor stability, turnover, and transcriptional activity ^32^ ^33^. Within the ROR subfamily, the hinge domain is the least conserved region, suggesting that it may confer receptor-specific regulatory functions rather than serving solely as a structural linker. Consistent with this idea, a two-residue (L92 and S93) substitution in the RORγ hinge selectively impairs ubiquitination of a lysine residue (K69) within the DBD without altering coactivator recruitment or DNA-binding activity. This finding demonstrates that the ROR hinge can allosterically regulate post-translational modification status at a structurally distant site within the receptor ^34^.

Here, we address a major gap in understanding RORβ regulation by demonstrating that the lysine acetyltransferase p300 directly acetylates RORβ, while SIRT1 mediates its deacetylation. Using RIME coupled with nano LC-tandem mass spectrometry, we defined the interactome of RORβ, and we mapped acetylation sites and assessed their interplay with ubiquitination and SUMOylation. Our findings reveal that p300 and SIRT1 coordinate a network of PTMs that govern RORβ stability, turnover, and transcriptional activity. Notably, an enzymatically inactive SIRT1 variant still partially stabilizes RORβ, uncovering a non-catalytic scaffolding role for SIRT1 in RORβ–p300 complex formation. Together, these results establish a mechanistic framework for RORβ regulation and more broadly highlight how combinatorial PTM networks regulate nuclear receptor function, providing potential strategies for therapeutic targeting.

## RESULTS

### RORβ directly binds to p300 and SIRT1, forming a functional complex that controls its acetylation

RORβ is expressed in cartilage, and its expression is altered in osteoarthritis, yet the receptor’s molecular function in chondrocytes remains undefined. To establish an unbiased view of the RORβ protein interaction network in this disease-relevant cell type, we performed proteomic profiling in TC28a2 human chondrocytes. To find putative coregulators of RORβ, we used a modified version of rapid immunoprecipitation of endogenous proteins method (RIME) ^35^, which captures protein-protein interactions in nuclei through in-cell formaldehyde crosslinking. Given that TC28a2 cells express low levels of RORβ, HA-tagged RORβ (isoform 1 throughout this study) was transiently transfected into TC28a2 human chondrocytes. Across the union of two independent experiments, a total of 206 proteins were enriched at log_2_ fold change ≥ 1.5 and p-value < 0.05. Gene ontology analysis of this interactome was dominated by terms associated with transcriptional regulation, including regulation of transcription by RNA polymerase II, DNA-templated transcription, and sequence-specific DNA binding, consistent with the expected nuclear receptor function of RORβ (**Fig. S1A**). Among the high-confidence interactors, the lysine acetyltransferase p300 (EP300) and the NAD⁺-dependent deacetylase SIRT1 emerged as candidates for post-translational regulation of RORβ (**Fig. 1A**). Triplicate measurements demonstrated reproducible enrichment of RORβ, p300, and SIRT1 over HA-peptide-blocked controls (**Fig. 1B**), which is expanded to show a heatmap of the top 50 interactors in **Fig. S1B**, indicating clear enrichment in the experimental samples over control across replicates.

**Figure 1.**
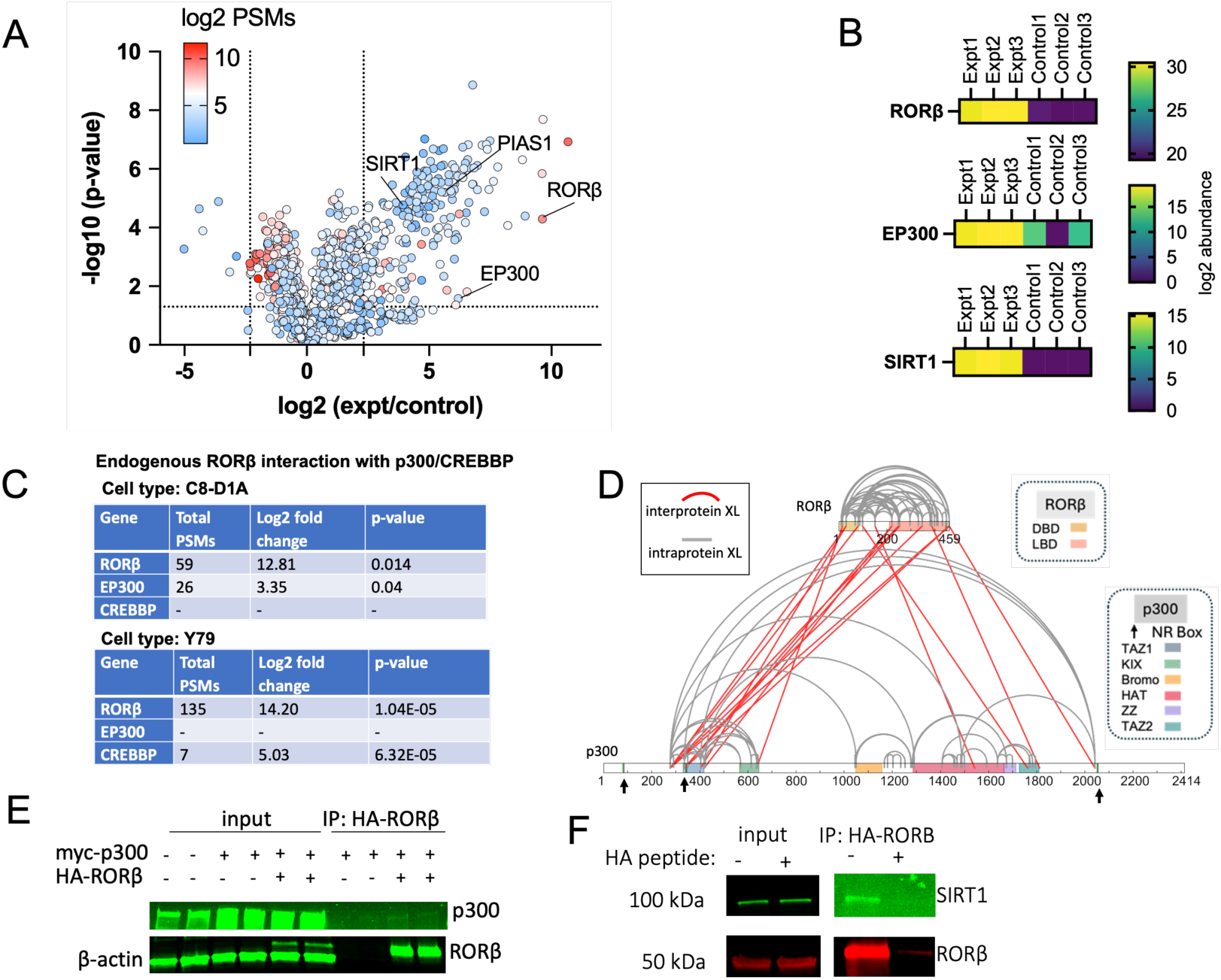
Proteomic profiling of the RORB interactome identifies p300/CREBBP and SIRT1 as key binding partners. (A) In-cell crosslinking followed by immunoprecipitation of transiently transfected HA-RORβ in TC28a2 cells. The volcano plot shows log2 fold change (experimental/control) on the x-axis and −log10(p-value) on the y-axis; point color indicates log2 total PSMs. Dotted lines denote the significance thresholds (log2 fold change > 2 and p ≤ 0.05. n = 3 biological replicates). (B) Heatmaps of protein abundance for RORβ, EP300, and SIRT1 across replicates from the experiment shown in (A). (C) Endogenous RORβ interaction with p300/CREBBP confirmed by immunoprecipitation mass spectrometry in two cell types with endogenous RORβ expression: C8-D1A mouse cerebellar astrocytes (RIME workflow, n=2) and Y79 human retinoblastoma cells (standard IP-MS workflow, n=3). (D) Crosslinking mass spectrometry of recombinant RORβ and p300 using DSSO crosslinker. Interprotein crosslinks (red) and intraprotein crosslinks (gray) are mapped onto the RORβ and p300 sequences. I Co-immunoprecipitation of RORβ and p300 in HEK293T cells. Cells were transfected with empty vector, HA-RORβ, or myc-p300 as indicated, and HA-RORβ was immunoprecipitated with an anti-HA antibody. Input and IP fractions were immunoblotted for p300 and RORβ. (F) Co-immunoprecipitation of RORβ and SIRT1 in HEK293T cells co-transfected with flag-SIRT1 and HA-RORβ. HA-RORβ was immunoprecipitated with an anti-HA antibody; excess HA peptide was used to block HA-RORβ binding to the anti-HA beads. Input and IP fractions were immunoblotted for SIRT1 and RORβ.

To determine whether the RORβ–p300/CBP and RORβ–SIRT1 associations occur without overexpression of receptor (endogenous expression levels), immunoprecipitation–mass spectrometry (IP-MS) was performed in C8-D1A mouse cerebellar astrocytes (RIME workflow with formaldehyde crosslinking) and Y79 human retinoblastoma cells (standard IP-MS without crosslinking), cell lines that robustly express RORβ. Experiments in both cell lines recovered the RORβ-p300/CBP interaction, with cell-type-specific paralog selectivity: C8-D1A cells yielded EP300 co-precipitation, while Y79 cells yielded CREBBP (**Fig. 1C**), indicating that RORβ engages both p300 paralogs in a cell-type-dependent manner.

To map the architecture of the RORβ–p300 interaction, recombinant RORβ and p300 were subjected to in vitro crosslinking with DSSO, followed by tandem mass spectrometric identification of crosslinked peptides using XlinkX in Proteome Discoverer. Multiple inter-protein crosslinks were detected. Two of the three NR-box motifs (LXXLL) in p300 crosslinked with K221, K253, and K459 of RORβ; and K2042 of p300, which is in proximity to the LQNLL NR-box motif (p300 residues 2051-2055), crosslinked with RORβ K459 (**Fig. 1D**). Additional inter-protein crosslinks mapped to the TAZ1 and HAT domains of p300, the latter through contact with RORβ-p300 interactions with both NR-box motifs and well-ordered coactivator domains. Validation of RORB-SIRT1 interaction was performed using immunoprecipitation of HA-RORβ from HEK293T cells followed by Western blotting confirming recovery of endogenous SIRT1, when in the absence of HA-peptide-blocking control (**Fig. 1F**). Similarly, reciprocal co-immunoprecipitation with myc-tagged p300 confirmed the RORβ-p300 interaction (**Fig. 1E**).

### p300-mediated acetylation of RORβ is reversed by SIRT1-dependent deacetylation

Determination of p300/CBP and SIRT1 as direct interactors with RORβ raised the possibility the receptor was controlled by opposing but coordinated acetyltransferase–deacetylase activities^31^. HA-RORβ was co-transfected into HEK293T cells with or without p300 and cells were treated with 50 µM EX-527 (a SIRT1-selective inhibitor), 1 mM nicotinamide (NAM; a broad sirtuin inhibitor), and 0.4 µM trichostatin A (TSA; a class I/II HDAC inhibitor), or DMSO. As determined by anti-acetyl lysine Western blotting, co-transfection with p300 yielded the largest magnitude of induction of RORβ acetylation when in the presence of EX-527; NAM yielded a smaller but measurable increase; whereas neither DMSO nor TSA conditions resulted in detection of p300-dependent acetylation (**Fig. 2A, C**). These findings indicate that the dominant deacetylase opposing p300 action on RORβ is a sirtuin, and not a class I/II HDAC. Co-transfection with p300 also lead to increased steady-state abundance of RORβ (**Fig. 2A, B**).

**Figure 2.**
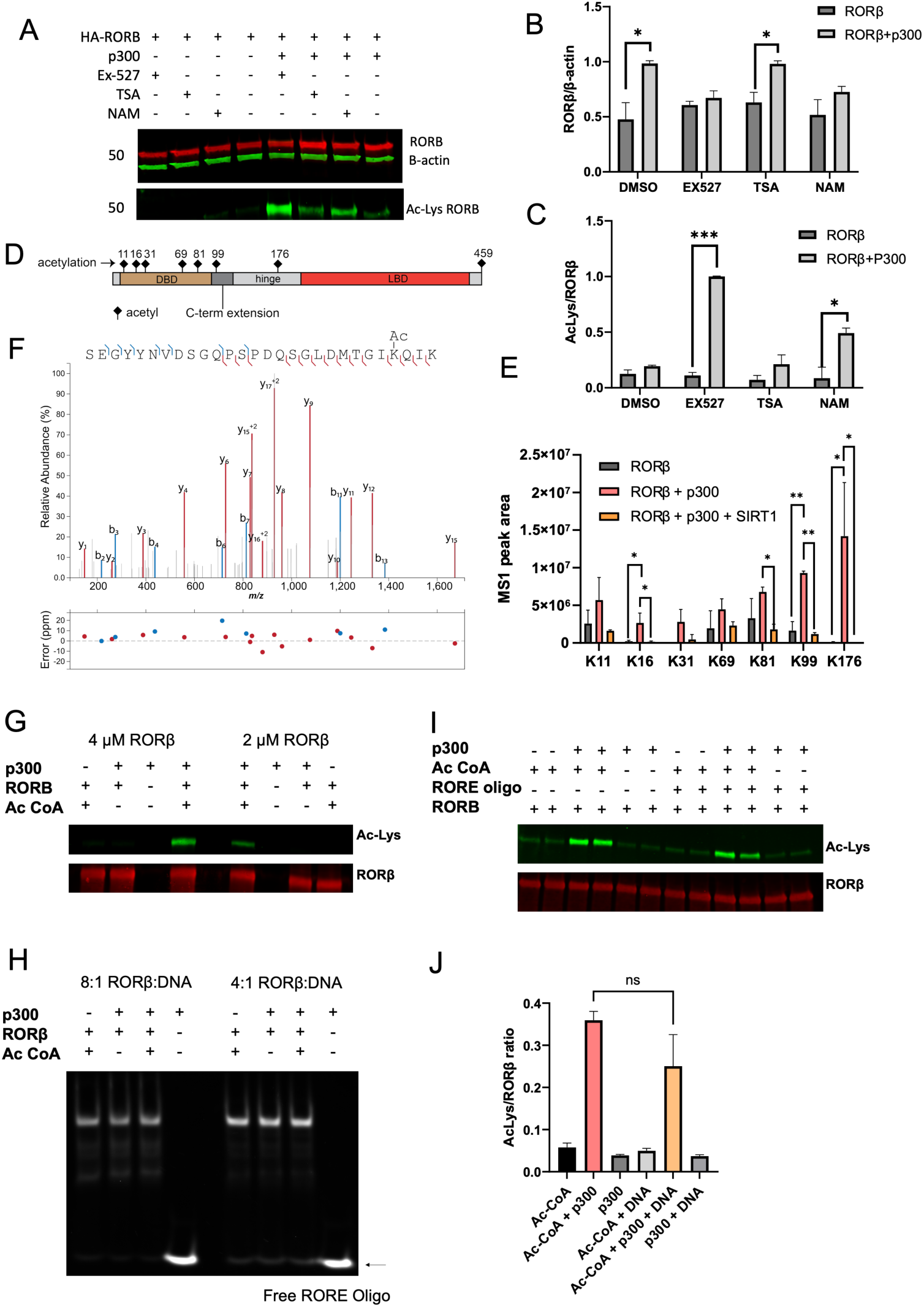
p300 acetylates RORβ at eight lysine site and does not alter DNA binding. (A) HEK293T cells were transiently transfected with HA-RORβ with or without p300 and treated with the indicated inhibitors: EX-527(50µM, SIRT1 inhibitor), TSA (400 nM, class I/II HDAC inhibitor), or NAM (1mM, broad sirtuin inhibitor). Lysates were immunoblotted for RORβ, β-actin (top), and acetyl-lysine on immunoprecipitated HA-RORβ (bottom). (B) Quantification of RORβ / β-actin ratio across conditions in (A) (n=3, two-way ANOVA). (C) Quantification of acetyl-lysine/ RORβ ratio across conditions in (A) (n=3, two-way ANOVA) (D) Schematic of RORβ domain architecture showing the positions of the eight acetylation sites identified by MS (K11, K16, K31, K69, K81, K99 in the DBD; K176 in the hinge; K459 in the LBD). I MS1 peak areas of acetylated peptides from HA- RORβ immunoprecipitated from HEK293T cells co-transfected with empty vector, p300, or p300+SIRT1 p300 co-transfection increases acetylation at all identified sites; SIRT1 co-transfection reduces acetylation at all sites (n=3). (F) Representative MS2 spectrum for the K176 acetylated RORβ peptide, with the mass error (ppm) for assigned fragment ions shown below. Spectrum annotated with IPSA (https://coonlabs.bmolchem.wisc.edu/). (G) Full-length recombinant RORβ was incubated with full-length recombinant p300 in the presence or absence of acetyl-CoA (1mM). Immunoblot for pan-acetyl-lysine confirms acetylation of RORβ only when both p300 and acetyl-CoA are present. (H) EMSA of acetylated or non-acetylated RORβ with a 40-bp RORE oligonucleotide at RORβ: DNA molar ratio of 8:1 and 4:1. No detectable difference in DNA binding is observed. (I) Full-length recombinant RORβ was incubated with full-length recombinant p300 and acetyl-CoA in the presence or absence of a RORE oligonucleotide, as indicated. Immunoblot of pan-acetyl-lysine (top) and total RORβ(bottom). (J) Quantification of acetyl-lysine/ RORβ ratio from the experiment in (I) (n=3; unpaired t-test, ns= not significant). Statistical comparisons: *p ≤ 0.05; **p ≤ 0.01; *** p ≤ 0.001; ****p ≤ 0.0001; ns, not significant. Error bars represent mean ± SEM).

To quantitate amino acid residue level acetylation, HA-RORβ was immunoprecipitated from HEK293T cells co-transfected with empty vector, p300, or p300 plus SIRT1, and analyzed by LC-MS/MS. Across two digestion strategies (trypsin alone, and trypsin combined with chymotrypsin), eight acetylation sites were identified: K11, K16, K31, K69, K81, and K99, all within the DBD; K176 within the hinge domain; and K459, which resides at the C-terminus of the LBD (**Fig. 2D**). Quantification of peptide MS1 peak areas using Skyline showed that p300 co-transfection significantly increased acetylation levels at most identified sites, while SIRT1 co-transfection reduced acetylation at all sites **(Fig. 2E**). Acetylation levels of RORB K176 were near the limits of detection in the presence of endogenous levels of p300/CBP, yet it was robustly induced by overexpression of p300, and levels were reduced back to basal by SIRT1 co-expression. The MS/MS spectrum supporting K176 acetylation assignment is shown in **Fig. 2F**.

To distinguish catalytic from non-catalytic SIRT1 effects on RORβ, LC-MS/MS was used to quantitate acetylation levels in the presence of wildtype SIRT1 and the catalytically inactive H363Y SIRT1 (**Fig. S2A**). H363Y SIRT1 failed to reduce RORβ acetylation levels: K176 acetylation with or without expression of H363Y were indistinguishable and substantially higher than with expression of wildtype SIRT1. This demonstrates that SIRT1 reverses RORβ acetylation through its deacetylase activity, and not by displacement of p300. Combined, these data identify RORβ as a p300 acetylation substrate, define eight acetyl-lysine sites concentrated in the DBD and at K176 in the hinge, and establish SIRT1 as the principal deacetylase opposing p300 via a catalytic mechanism. K176 emerged as the most SIRT1-sensitive site and was prioritized for further mechanistic characterization.

### p300-mediated acetylation does not alter RORβ DNA binding

Six of the eight RORβ acetylation sites identified as shown in **Fig. 2** are within the DBD, raising a question of whether acetylation alters DNA binding. Full-length recombinant RORβ was incubated with active full-length recombinant p300 in the presence or absence of the acetyl-CoA. Immunoblotting for pan-acetyl-lysine confirmed acetylation of RORβ only when both p300 and acetyl-CoA were present (**Fig. 2G**). The ability of acetylated RORβ to bind DNA was examined with an electrophoretic mobility shift assay (EMSA) using a 40-bp RORE oligonucleotide at molar ratios of RORβ to DNA of 8:1 and 4:1. Acetylation did not alter RORβ–DNA binding, as no differences were observed in either the shifted complex or free DNA at either stoichiometry (**Fig. 2H**). To test whether RORβ binding to RORE DNA would interfere with p300-mediated acetylation, receptor was incubated with p300 and acetyl-CoA in the presence and absence of the RORE oligonucleotide. No significant difference in acetyl-lysine/RORβ ratio was observed (**Fig. 2I, J**). Together, these data establish that p300-mediated acetylation does not alter receptor binding to DNA, and DNA binding does not alter acetylation of RORβ.

### p300 and SIRT1 regulate RORβ stability and transcriptional output via K176

In order to evaluate the impact of p300 and SIRT1 on receptor stability and transcriptional activity, HEK293T cells were co-transfected with varying combinations of HA-RORβ, myc-p300, flag-SIRT1 wild-type (WT), or flag-SIRT1-H363Y, as indicated in **Fig. 3A**. RORβ abundance was significantly increased by p300 co-transfection and further elevated by the combination of p300 and SIRT1 WT, which produced the highest RORβ/β-actin ratio of all conditions tested (p < 0.05, **Fig. 3A, B**). Co-transfection of p300 with the catalytically inactive SIRT1 H363Y mutant resulted in less stabilization of RORβ than wild-type SIRT1, indicating that SIRT1 deacetylase activity contributes to RORβ stability. A cycloheximide (CHX) chase experiment performed 48 h post-transfection showed that RORβ half-life was most significantly extended by the combination of p300 and SIRT1 WT (**Fig. 3C**). In contrast, p300 alone, SIRT1-H363Y alone, and the p300+SIRT1-H363Y all resulted in rapid RORβ decay comparable to RORβ alone. The partial stabilization observed with SIRT1-H363Y in the presence of p300 suggests a scaffolding role for SIRT1 in a RORβ-p300-SIRT1 complex, independent of its catalytic activity. Reporter assays using a 5X RORE firefly luciferase construct showed modest activation with p300 (∼2.5-fold over RORβ alone), greater activation with SIRT1 (∼5-fold), and marked increase with the combination of p300 and SIRT1 WT (∼16-fold) (**Fig. 3D**). This pattern paralleled steady-state RORβ abundance, indicating that transcriptional output is tightly coupled to receptor levels.

**Figure 3.**
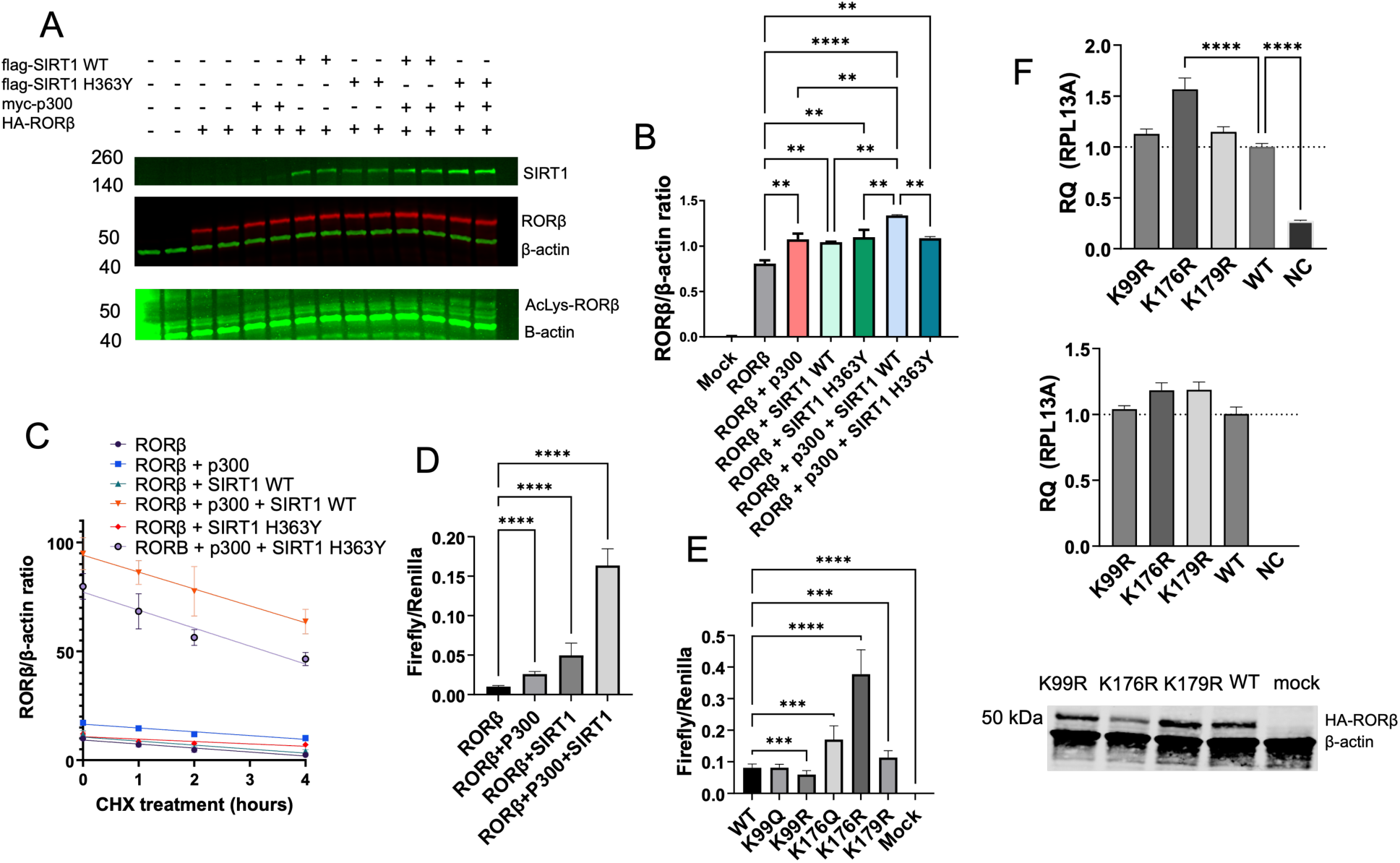
p300 and SIRT1 regulate RORβ stability and transcriptional activity. (A, B) RORβ protein abundance under co-transfection conditions. HEK293T cells were transfected with HA-RORβ in combination with myc-p300, flag-SIRT1 WT, or the catalytically inactive flag-SIRT1-H363Y mutant as indicated and lysates were immunoblotted for SIRT1, RORβ, β-actin, and acetyl-lysine RORβ (A). RORβ abundance (RORβ/β-actin) is quantified in (B). (n = 2; one-way ANOVA). (C) RORβ protein half-life determined by cycloheximide (CHX) chase across the same co-transfection conditions. RORβ/β-actin is plotted against time of CHX treatment (n = 2). (D) Transcriptional activity of RORβ measured by RORE-driven luciferase reporter assay across the same co-transfection conditions. Reporter activity (firefly/Renilla) increases progressively with p300, SIRT1, and their combination (n = 5; one-way ANOVA). (E) RORE-driven reporter activity of HA-RORβ acetylation-site mutants and mock control in HEK293T cells (n = 4; one-way ANOVA). (F) Effect of RORβ acetylation-site mutations on target gene expression in MG63 cells. Cells were transiently transfected with HA-tagged WT RORβ or the K99R, K176R, or K179R mutants. mRNA levels of the RORβ target gene NR1D1 (top) and of RORβ itself (middle, transfection control) were measured by qPCR. RORβ WT and mutant protein levels were assessed by anti-HA immunoblot with β-actin as a loading control (bottom). The K176R mutant produces the highest induction of NR1D1 despite comparable or lower protein expression than WT, consistent with elevated transcriptional activity. (n = 3, one-way ANOVA).

To identify the specific acetylated lysines responsible for these effects, RORβ mutants were generated at K99 and K176, substituting lysine with arginine (R, mimicking the non-acetylated state) or glutamine (Q, partially mimicking the acetylated state). K179R was also generated, as K179 resides within a canonical SUMOylation site (discussed below). RORE-driven reporter activity was measured for each mutant (**Fig. 3E**). K99Q and K99R exhibited reporter activity comparable to WT, while K179R showed a modest increase relative to WT. In contrast, K176Q displayed approximately twofold higher activity than WT, whereas K176R produced the strongest effect (∼5-fold). Subcellular fractionation confirmed that these differences were not due to altered nuclear accumulation of the mutants (**Fig. S2B**).

To validate the K176R phenotype on an endogenous RORβ target gene, WT, K99R, K176R, and K179R constructs were expressed in MG63 cells, and NR1D1 expression was measured by qPCR (**Fig. 3F**). K176R produced significantly higher NR1D1 expression than WT, whereas K99R and K179R showed comparable levels. Notably, K176R appeared modestly lower in abundance than WT despite similar or slightly elevated target gene induction, suggesting increased transcriptional activity per molecule. Together, these data establish that K176 is the primary determinant of RORβ activity among the sites tested.

### RORβ is SUMOylated at K179, and K176 acetylation primes UBC9-mediated SUMOylation

Sequence analysis of RORβ identified a highly conserved canonical (ψKxE/D) SUMOylation motif at K179, located in close proximately to the p300/SIRT1 target residue K176, raising the possibility of acetylation-SUMOylation crosstalk in the RORβ hinge region (**Fig. S3A, B**). In vitro SUMOylation assays using full-length recombinant RORβ, SUMO E1 activating enzyme, UBC9 E2 conjugating enzyme, and ATP demonstrated conjugation of SUMO2 and SUMO3 to RORβ in an ATP-dependent manner, producing a ∼100 kDa modified species (**Fig. S4A**). No appreciable SUMO1 conjugation was detected, despite robust SUMO1 modification of the RanGAP1 positive control (**Fig. S3C**). Mass spectrometry confirmed K179 as the SUMO acceptor site, with the characteristic QQQTGG SUMO2 remnant attached to K179 following trypsin/chymotrypsin digestion (**Fig. S4B**).

To investigate whether K179 serves as the SUMO acceptor site in cells, HA-RORβ WT or the K179R mutant was co-transfected into HEK293T cells with Flag-SUMO2 and UBC9, with or without PIAS1, which was identified as a RORβ interactor in the RIME interactome (**Fig. 1, Fig. S4C**). A SUMO-modified RORβ species at ∼100 kDa was present in WT cells co-expressing SUMO2 and UBC9 and was strongly enhanced by PIAS1 (**Fig. 4A**). In contrast, the K179R mutant produced essentially no SUMO-modified RORβ under all conditions tested, indicating that K179 is the primary SUMO acceptor site on RORβ in cells. Independent mass spectrometric identification of the K179-modified peptide from HA-RORβ co-transfected with UBC9, SUMO2, and PIAS1 confirmed K179 as the SUMOylation site in the cellular context (**Fig. 4B**).

**Figure 4.**
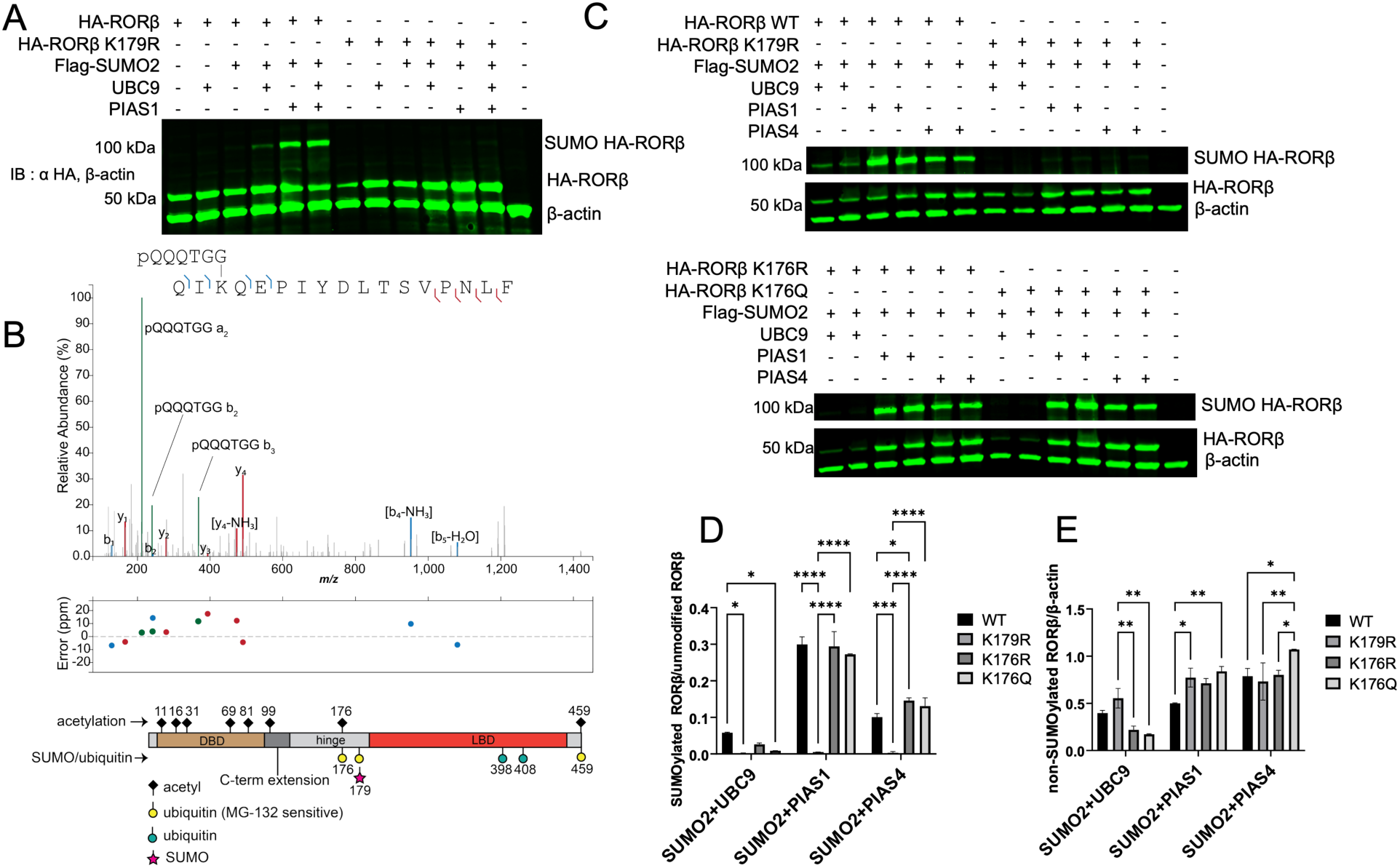
RORB is SUMOylated at K179, and K176 acetylation primes SUMOylation. (A) Western blot of SUMOylation of HA-RORβ WT and the K179R mutant in HEK293T cells transfected with HA-RORβ (WT or K179R), Flag-SUMO2, UBC9, and PIAS1 as indicated. Lysates were immunoblotted with anti-HA (detecting SUMO-HA-RORβ at ∼100 kDa and unmodified HA-RORβ at ∼50 kDa) and β-actin. (B) Confirmation of K179 as a SUMOylation site in RORβ by IP-MS of HA-RORβ from HEK293T cells co-transfected with UBC9, SUMO2, and PIAS1. The representative MS2 spectrum of the K179-containing peptide was obtained following trypsin/chymotrypsin digestion; diagnostic fragment ions from the QQQTGG SUMO2 remnant are shown in green, with the mass error (ppm) for assigned fragment ions shown below. Spectrum annotated with IPSA. The schematic at bottom summarizes the acetylation, ubiquitination, and SUMOylation sites mapped on RORβ across this study. (C) SUMOylation of HA-RORβ WT and K179R (top) and of K176R and K176Q (bottom) in HEK293T cells, transfected with Flag-SUMO2 and UBC9, with PIAS1 or PIAS4 as indicated. Lysates were immunoblotted with anti-HA (SUMO-HA-RORβ at ∼100 kDa, HA-RORβ at ∼50 kDa) and β-actin. (D, E) Quantification of the immunoblots in (C). (D) Normalized abundance of non-SUMOylated RORβ. (E) Ratio of SUMOylated to non-SUMOylated RORβ (n = 2 biological replicates; two-way ANOVA).

To test whether K176 is mechanistically coupled to K179 SUMOylation, SUMOylation of the K176R (non-acetylatable) and K176Q (acetylation mimic) mutants was compared with WT in the presence of SUMO2 and UBC9, with or without PIAS1 or PIAS4 (**Fig. 4C**). In the absence of exogenous E3 ligase, both K176R and K176Q exhibited substantially reduced SUMOylation relative to WT, with the SUMOylated to non-SUMOylated ratio reduced to approximately one-fifth of the WT level (**Fig. 4E**, leftmost group). In contrast, co-expression of PIAS1 or PIAS4 resulted in robust SUMOylation of WT, K176R, and K176Q at comparable levels, restoring SUMOylation of the K176 mutants to WT-equivalent efficiency (**Fig, 4C, 4E**). Together, these results identify K179 as the SUMO acceptor site of RORβ, demonstrate preference for SUMO2/3, and establish K176 acetylation as a priming step for UBC9-mediated SUMOylation at K179 that is bypassed by PIAS1 and PIAS4.

### SIRT1 inhibition enhances PIAS1-mediated SUMOylation of RORβ

To test whether acetylation-SUMOylation crosstalk operates dynamically in cells, HEK293T cells co-transfected with HA-RORβ (WT or K179R), p300, and PIAS1 were treated with the SIRT1 inhibitor EX-527 (50 µM) for 18 h prior to harvest. Acetyl-lysine RORβ, SUMOylated RORβ, and total RORβ were detected by multiplexed immunoblot (**Fig. 5A, 5B**). In WT-expressing cells, acetyl-lysine RORβ was induced by p300 and further elevated by EX-527, consistent with SIRT1 acting as the rate-limiting deacetylase for K176 (**Fig. 5A**). SUMOylated RORβ (∼100 kDa) was detected only upon PIAS1 co-expression and was further enhanced by EX-527. In contrast, no SUMO-modified RORβ was detected in K179R-expressing cells under any condition (**Fig. 5B**), confirming that K179 is required for SUMOylation in this context.

**Figure 5.**
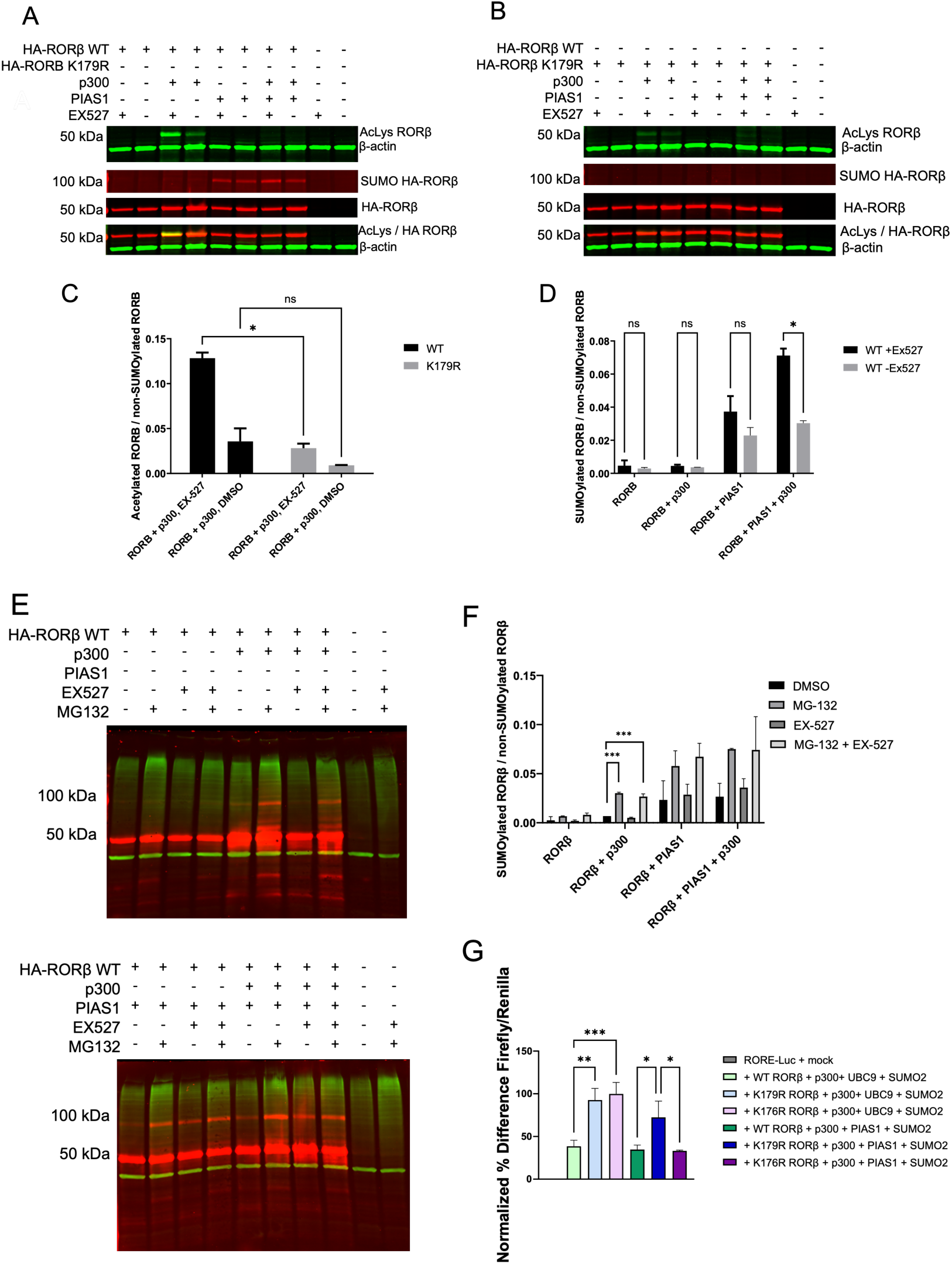
SIRT1 inhibition promotes PIAS1-dependent K179 SUMOylation that targets RORβ for proteasomal degradation and represses transcriptional activity. (A) HEK293T cells were co-transfected with HA- RORβ (WT), P300 and PIAS1 as indicated and treated with EX-527(50µM, SIRT1 inhibitor) for 18h prior to harvest. Acetyla-lysine RORβ, SUMOylated RORβ, and total RORβ were detected by multiplexed immunoblot. AcLys- RORβ was strongly induced by p300 and further elevated by EX-527, consistent with SIRT1 as the rate-limiting deacetylase for K176. (B) As in (A), using HA- RORβWT or the SUMO-deficient K179 mutant. SUMOylated RORβ (∼100kDa) was detected a only when PIAS1 was co-expressed and was enhanced by EX- 527 in WT-expressing cells; no SUMO- RORβ band was detected in K179R-exoressing cells under any condition. (C) Quantification of acetylated RORβ relative to non-SUMOylated RORβ for WT and K179R (n = 2; two-tailed t-test). (D) Quantification of SUMOylated RORβ relative to non-SUMOylated RORβ for WT RORβ in the presence or absence of EX-527 across the indicated conditions (n = 2; two-tailed t-test). (E) HEK293T cells were transiently transfected with combinations of HA-RORβ, p300, and PIAS1 as indicated, and treated with the SIRT1 inhibitor EX-527 and/or the proteasome inhibitor MG-132. Lysates were immunoblotted with anti-HA (top: detecting SUMO-HA-RORβ at ∼100 kDa and unmodified HA-RORβ at ∼50 kDa) and anti-ubiquitin (bottom). (F) Quantification of the ratio of SUMOylated to non-SUMOylated RORβ (n = 2; two-way ANOVA). (G) RORE-driven luciferase reporter activity of WT RORβ and mutants in HEK293T cells under two SUMOylation-driving conditions, p300 + UBC9 + SUMO2 and p300 + PIAS1 + SUMO2, normalized to the RORE-Luc + mock control (n = 4; one-way ANOVA without correction for multiple comparisons).

Quantification confirmed that EX-527 significantly increased acetylation in WT-expressing cells co-transfected with p300 (**Fig. 5C**). Analysis of the SUMOylated-to-non-SUMOylated RORβ ratio across four conditions (RORβ alone, +p300, +PIAS1, +p300+PIAS1), with or without EX-527, revealed a stepwise increase in SUMOylation, with maximum levels observed in cells co-expressing both p300 and PIAS1 in the presence of EX-527 (**Fig. 5D**). Of note, only the RORβ+p300+PIAS1 condition showed a statistically significant EX-527-dependent increase, with SUMOylation approximately doubling relative to DMSO alone. Together, these results indicate that SIRT1 activity is rate-limiting for SUMOylation when both acetylation machinery and an E3 ligase are present, supporting a model in which K176 acetylation primes K179 SUMOylation in a dynamic, deacetylase-regulated process manner.

### SUMOylation targets RORβ for proteasomal degradation and reduces transcriptional activity

HEK293T cells expressing HA-RORβ were treated with MG-132 (20 µM; proteasome inhibitor) in combination with p300 co-expression, EX-527 (30 µM) or both. A band corresponding to SUMOylated RORβ (∼100 kDa) accumulated specifically upon p300 co-expression and MG-132 treatment in cells lacking exogenous SUMO machinery, indicating that endogenous SUMO machinery modifies RORβ and that the resulting SUMOylated pool is normally cleared by the proteasome (**Fig. 5E, top**). Co-expression of PIAS1 produced robust SUMOylated RORβ, which was further enhanced by MG-132 (**Fig. 5E, bottom**). Quantitative analysis identified MG-132 as the dominant factor impacting steady-state SUMOylated RORβ levels, suggesting that proteasomal turnover, rather than rate of SUMO conjugation, is the primary determinant of steady-state SUMOylated RORβ accumulation when both p300 and an E3 ligase are present (**Fig. 5F**).

RORE-driven reporter assays in HEK293T cells were used to compare WT, K179R, and K176R RORβ under two SUMOylation-promoting conditions: p300+UBC9+SUMO2 (E3-independent), in which K176 acetylation primes UBC9-mediated SUMOylation at K179, and p300+PIAS1+SUMO2, in which PIAS1 bypasses the priming requirement. As shown in **Fig. 5G**, under p300+UBC9+SUMO2 conditions, both K179R and K176R mutants exhibited increased reporter activity, approximately 2.5- and 2.8-fold higher than WT, respectively. In contrast, under p300+PIAS1+SUMO2 conditions, K179R activity remained elevated, whereas K176R activity returned to WT levels, consistent with PIAS1 restoring K179 SUMOylation of the K176R mutant. Across both conditions, K179R functions as a constitutive non-SUMOylatable control, exhibiting elevated activity regardless of E3 ligase identity, thereby confirming that loss of K179 SUMOylation derepresses RORβ activity. In contrast, K176R acts as a sensor of the K176 priming requirement: its activity is elevated when SUMOylation depends on K176 acetylation priming (UBC9 alone) but is restored to WT levels when PIAS1 bypasses this requirement and directly SUMOylates K179.

### p300 displaces UBR5 from RORβ, and SIRT1 activation reduces PIAS1 association

IP-MS was performed on HA-RORβ from HEK293T cells across four co-transfection conditions: RORβ alone (R), RORβ+p300 (RP), RORβ+SIRT1 (RS), and RORβ+p300+SIRT1 (RSP). The RSP condition yielded the highest RORβ abundance based on peptide spectral match (PSM) counts (**Fig. 6B**), consistent with the elevated RORβ levels observed by Western blot under this condition (**Fig. 3**). The E3 ubiquitin ligase UBR5 was the most abundant RORβ interactor by PSM count and showed substantially reduced association in the RP and RSP conditions compared to R and RS when normalized to RORβ PSMs (**Fig. 6C**). These data indicate that p300 co-expression promotes displacement of UBR5 from RORβ.

**Figure 6.**
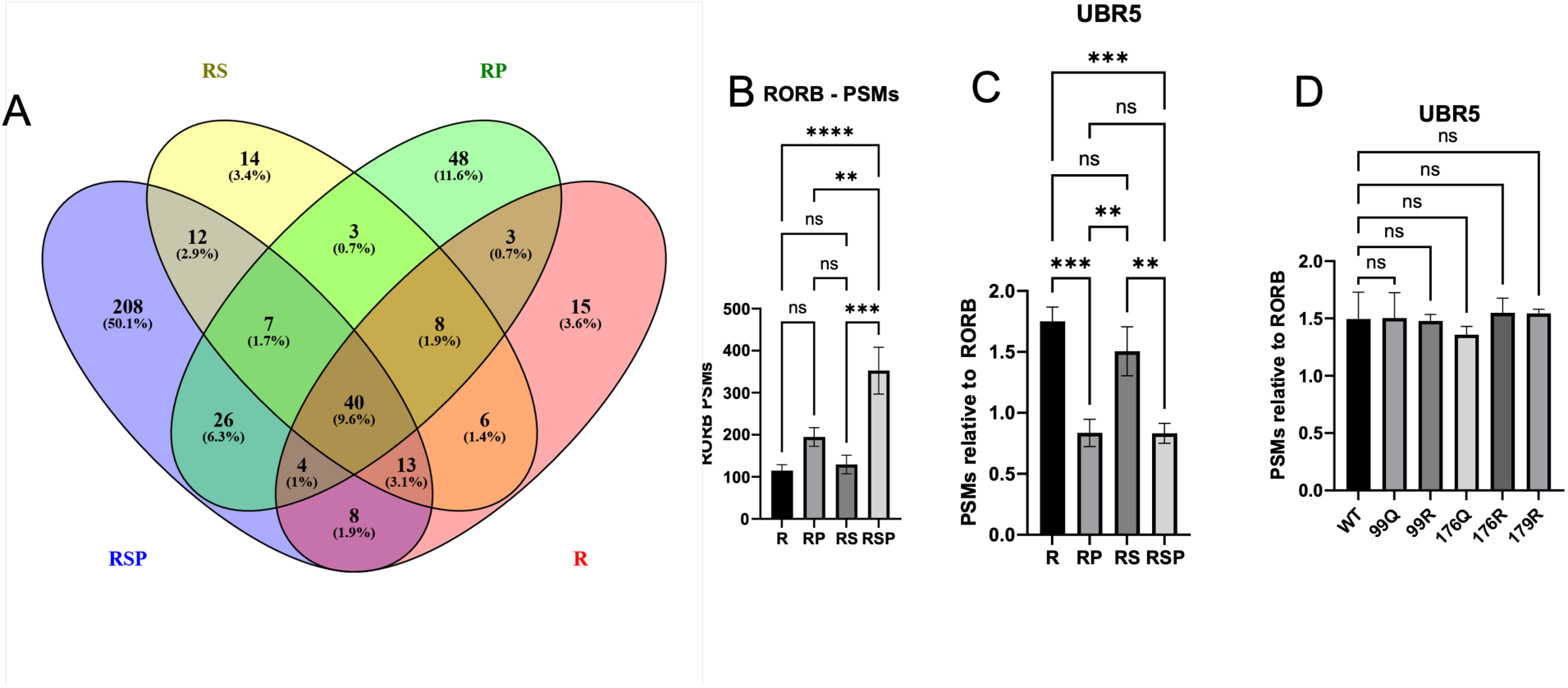
Acetylation state regulates RORβ association with ubiquitin ligases and SUMO machinery. (A) Four-way comparison of RORβ interactors identified by anti-HA IP-MS of HA-RORβ from HEK293T cells transfected with RORβ alone (R), RORβ + p300 (RP), RORβ + SIRT1 (RS), or RORβ + p300 + SIRT1 (RSP). 40 proteins were detected as core interactors across all four conditions, among which UBR5 was most abundant by PSM count. (B) RORβ peptide-spectrum matches (PSMs) recovered across the four conditions. RORβ recovery is highest in the RSP condition (n = 3; one-way ANOVA). (C) UBR5 PSMs normalized to RORβ PSMs across the four conditions. Co-expression of p300 (RP and RSP) reduces the association of UBR5 with RORβ (n = 3; one-way ANOVA). (D) UBR5 PSMs normalized to RORβ PSMs for HA-RORβ WT and the mutants, transfected without exogenous p300 (n = 2; one-way ANOVA). (E) Volcano plot of RORβ interactors from RIME of HA-RORβ in TC28a2 chondrocytes treated with the SIRT1 activator SRT2104 versus DMSO. The x-axis shows log2 fold change (SRT2104/DMSO) and the y-axis −log10(p-value); dotted horizontal line denotes threshold of p ≤ 0.05. PIAS1 is labeled and is significantly reduced upon SIRT1 activation (n = 3).

To test whether K-site mutations alter UBR5-RORβ association, IP-MS was performed on WT, K99Q, K99R, K176Q, K176R, and K179R RORβ under endogenous p300/CBP conditions (**Fig. 6D**). UBR5 association, normalized to RORβ, was comparable across all mutants, indicating that K99, K176, and K179 are not required for UBR5 binding under basal acetylation conditions. To examine the reciprocal effect of SIRT1 activity on RORβ-PIAS1 association, RIME-based IP-MS was performed on HA-RORβ in TC28a2 chondrocytes treated with the SIRT1 activator SRT2104 or DMSO alone. PIAS1 association was significantly reduced in RORβ immunoprecipitates from SRT2104-treated cells (log_2_ fold change ∼-0.4, p < 0.05; **Fig 6E**), consistent with SIRT1-mediated deacetylation at K176 reducing PIAS1 recruitment. Together, these results indicate that RORβ engages distinct ubiquitin-pathway proteins depending on its acetylation state: UBR5 associates with RORβ under endogenous acetylation levels and is displaced upon p300 co-expression, whereas PIAS1 is recruited under conditions that favor K176 acetylation and is reciprocally reduced when SIRT1 is pharmacologically activated.

### K176/K179 hinge mutations reshape the conformational landscape of the RORβ-p300 complex

To determine whether substitution of K176 or K179 alters the conformation of the RORβ-p300 complex, full-length recombinant RORβ (WT, K176R, or K179R) was incubated with active full-length recombinant p300 and a RORE DNA oligonucleotide, followed by crosslinking with the MS-cleavable crosslinker DSSO. Crosslink abundances were then compared between WT and each mutant for both intra- and inter-protein crosslinks (n=3) (**Fig. 7A**). In each comparison, a subset of crosslinks exhibited significant changes in abundance (p<0.05|log_2_FC|>1; **Fig. 7B**). Notably, these changes were predominantly observed in intramolecular p300 crosslinks, particularly within the HAT domain and the C-terminal TAZ2 region, rather than in RORβ intramolecular or RORβ-p300 intermolecular crosslinks. This pattern was consistent across both hinge mutants (K176R and K179R). These findings indicate that loss of lysines 176 or 179 reshapes p300 conformation within the RORβ-300-DNA complex, suggesting that the RORβ hinge region functions not merely as a local PTM site, but as an allosteric module that influences p300 structure.

**Figure 7.**
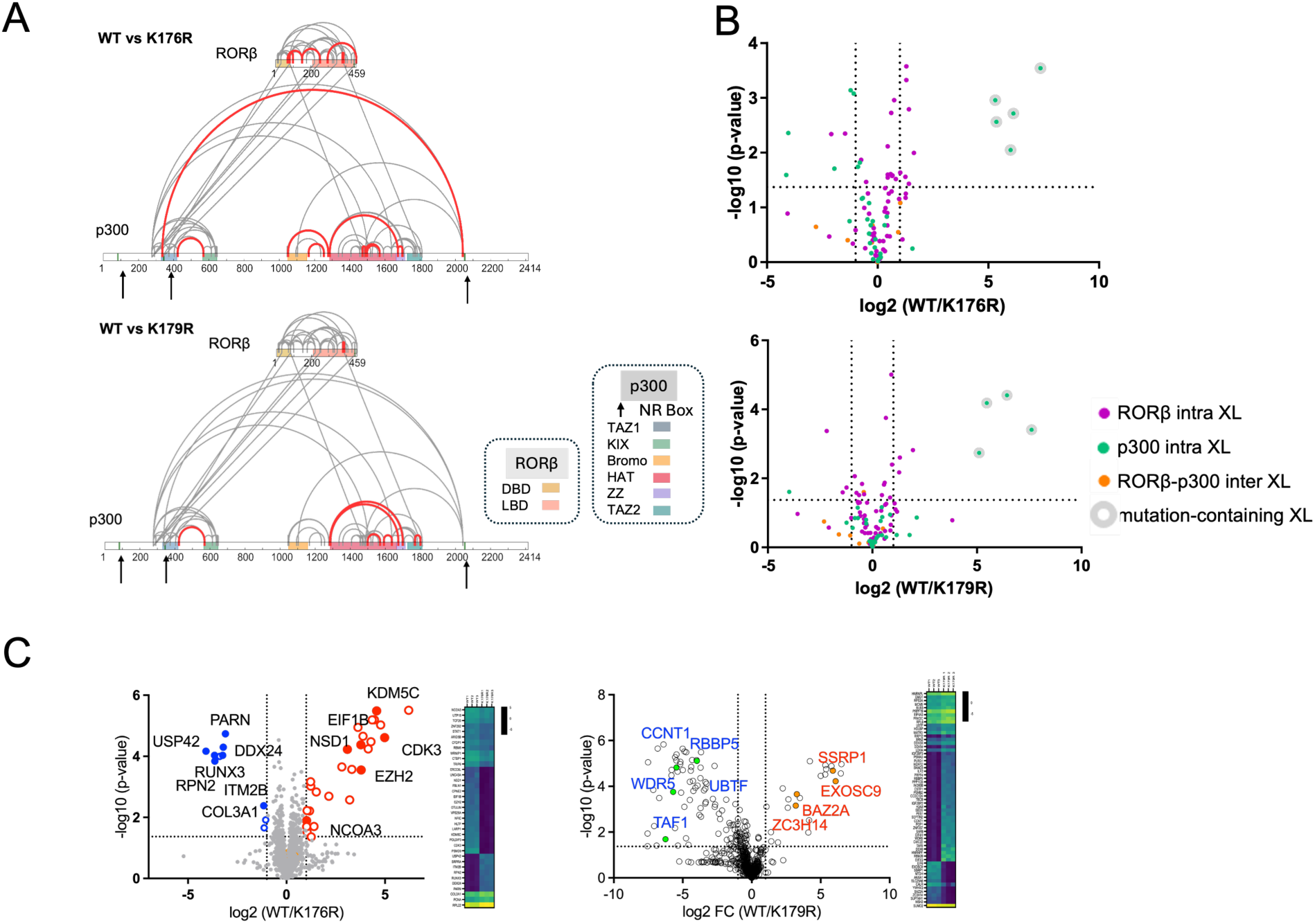
Quantitative XL-MS and RIME reveal conformational and coregulatory changes in p300-RORβ complexes responsive to K176 and K179 mutations. (A) Crosslink maps for full-length recombinant RORβ WT vs. K176 and WT vs. K179R mutants, incubated with p300 and RORE DNA oligomer and crosslinked with the MS-cleavable crosslinker DSSO. Crosslinks containing peptides spanning the RORβ mutation site are highlighted, as such crosslinks can only be detected in the WT sample (imputation prevents these crosslinks from showing an infinite fold change). Crosslinks with significantly changing abundance (p-value < 0.05, log_2_FC >|1|; n=3) are highlighted in red. RORβ and p300 domain architecture is annotated below each linear map (legend, right); arrow mark NR box motifs in p300. (B) Volcano plots showing the quantitative comparisons in (A) for WT vs 176R (top) and WT vs. K179R (bottom). Each point represents a unique crosslink, colored by category: RORβ intra-protein XL (purple), p300 intra-protein XL (green), RORβ-p300 inter-protein XL (orange), and mutation-containing XL (gray outline). The horizontal dotted line marks -log_10_(p-value) =0.05; vertical dotted lines mark log_2_FC = ± 1. (C) RIME comparing K176R or K179R to WT RORβ in TC28a2 chondrocytes (HA- RORβ IP, LFQ quantification). Left: K176R loses NR coactivation factors (blue) and gain DNA damage machinery (Red) relative to WT. Right: under conditions enhancing WT RORβ SUMOylation (SUMO2/PIAS1 co-transfection), K179R preferentially associates with transcriptional activation complexes (blue), while SUMOylated WT associates with transcriptional repressors and RNA surveillance factors (red/ orange). The non-overlapping interactor shifts indicate that K176 acetylation and K179 SUMOylation independently direct distinct coregulatory outputs (n=3).

To investigate whether this PTM-dependent conformational change translates into altered coregulator engagement in cells, RIME was performed in TC28a2 chondrocytes expressing HA-tagged WT, K176R, or K179R RORβ (**Fig. 7C**). Relative to WT, K176R showed reduced association with NR coactivators and increased association with DNA damage-related factors. In contrast, K179R, under conditions that enhance SUMOylation of RORβ WT, preferentially retained transcriptional activation complexes that were depleted from SUMOylated WT, which instead favored association with transcriptional repressors and RNA surveillance factors. The largely non-overlapping nature of these interactor shifts indicates that K176 acetylation and K179 SUMOylation independently gate access to distinct coregulatory complexes, providing a functional correlation to the conformational changes detected by XL-MS.

## DISCUSSION

RORβ is a transcription factor with critical functions in the central nervous system, retina, and bone, where even subtle changes in receptor abundance can have significant physiological consequences. While the receptor is enriched in retinal cells and in astrocytes, reduced RORβ expression has been linked to blindness, neurological disorders, osteoarthritis, and cancer. Despite these important roles, the post-translational mechanism that regulates RORβ abundance and activity has not been defined. Using IP-MS, crosslinking-MS, and targeted PTM mapping, we identify a coordinated acetylation-SUMOylation circuit centered on hinge residue K176 that links the acetyltransferase p300 and the deacetylase SIRT1 to two opposing RORβ fates; a stabilized, transcriptionally active state, and a SUMOylated state that is degraded via the proteasome and transcriptionally repressed.

p300 acetylates RORβ at six lysines within the DBD and at hinge residue K176, which is distinguished by being nearly undetectable at basal p300/CBP levels and almost completely reversed by SIRT1. This architecture, p300-dependent acetylation opposed by SIRT1 (validated using catalytically inactive H363Y SIRT1 and rescued by inhibition with EX-527), was previously established for the closely related paralog RORγ, where SIRT1 deacetylates the homologous DBD lysines (K69, K81, K99) to enhance transcriptional output ^36^. The present findings extend this acetylation/deacetylation logic from RORγ to RORβ but reveal key mechanistic distinction: the K176/K179 hinge module has no counterpart in the RORγ mechanism and represents the central advance of this work. Specifically, this module introduces a second, hinge-localized acetylation event that is mechanistically coupled to a downstream SUMOylation step, rather than acting solely through effects on DNA binding or coactivator recruitment.

A central and unexpected finding is that SIRT1 stabilizes RORβ through two distinct mechanisms. The catalytically active enzyme deacetylates K176 and, in combination with p300, produces the greatest extension of RORβ half-life and the highest RORE-reporter activity observed. In contrast, the catalytically inactive SIRT1 H363Y mutant still confers partial stabilization when co-expressed with p300, despite leaving K176 acetylation unchanged, indicating that SIRT1 can promote RORβ stability independently of its enzymatic activity, likely through a scaffolding function within a RORβ-p300-SIRT1 complex. This scaffolding role is supported by crosslinking-MS data showing that p300 engages RORβ through multiple structured contact points rather than a single interface. The RORβ DBD/LBD region crosslinks to two NR-box-proximal sites on p300, one adjacent to the TAZ1 domain (K336, near the LVLLL motif at residues 342-346) and another near the CREB-binding region (K2042, adjacent to the LQNLL motif at residues 2051-2055), along with an additional contact between the RORβ hinge (K151) and the p300 HAT domain. Notably, this hinge-HAT crosslink was detected in only one of two independent crosslinking experiments, one performed in the presence of RORE DNA and the other in its absence. This multivalent interaction suggests that RORβ is anchored to p300 through several structured interfaces simultaneously, potentially positioning the HAT domain in proximity to the hinge for acetylation of K176. Although we have not directly tested whether this docking configuration regulates p300 catalytic activity, these data support a model in which scaffolding interactions contribute to RORβ stabilization and coordinate enzymatic access to the hinge region.

This non-catalytic contribution of SIRT1 to nuclear receptor stability is not without precedent. SIRT1 has been reported to enhance glucocorticoid receptor (GR) transcriptional output independently of its deacetylase activity, acting instead as a structural component of the receptor’s coactivator complex, an effect that persists with the same catalytically inactive H363Y mutant used here and is similarly insensitive to EX-527 ^37^. The recurrence of this behavior across two structurally distinct nuclear receptors raises the possibility that SIRT1 functions more broadly as a scaffold within nuclear receptor transcriptional complexes, with its deacetylase activity providing an additional, receptor-specific layer of regulation rather than constituting its sole mode of action.

K176 acetylation functions as a priming signal for SUMOylation at the neighboring canonical SUMO site, K179. In the absence of an E3 ligase, UBC9-mediated SUMO conjugation occurs on K176 acetylated RORβ but not on the glutamine-substituted mimic, indicating that the acetyl modification itself, rather than the charge it confers, is required to support conjugation. PIAS1 and PIAS4 bypass this requirement, restoring SUMOylation to wild-type levels on both K176R and K176Q. Consistent with this model, SIRT1 inhibition increases both K176 acetylation and PIAS1-dependent SUMOylation in parallel. SUMOylated RORβ accumulates upon proteasome inhibition and is associated with reduced RORE-driven activity, placing K179 SUMOylation downstream of K176 acetylation in a pathway that attenuates, rather than enhances, RORβ output. Rather than functioning as a discrete on/off switch, the K176-Ac/K179-SUMO module operates as a continuous control point: SIRT1 activity determines the size of the acetylated, SUMO-competent RORβ pool, while the availability of an E3 ligase such as PIAS1 determines the efficiency with which this pool is converted into a degradation-prone, transcriptionally repressed species.

UBR5 was identified as the principal RORβ-associated E3 ligase in ubiquitination site mapping performed in HEK293T cells with co-expression of RORβ, p300, and SIRT1. Under conditions favoring p300-SIRT1 complex formation and maximal RORβ abundance, UBR5 association was reduced. UBR5 has recently been structurally characterized as a HECT-family ligase that primarily functions as a chain elongator, extending pre-ubiquitinated substrates rather than initiating ubiquitination de novo ^38,39^. This raises the possibility that the reduced UBR5 association observed here reflects altered substrate priming of RORβ, rather than simple steric displacement by p300, a distinction that would require direct assessment of RORβ ubiquitination prior to acetylation. The functional consequence of this regulatory circuit was confirmed at an endogenous RORβ target gene. When expressed at matched mRNA and protein levels in MG63 cells, K176R produced significantly higher NR1D1 expression than wild-type RORβ, whereas K99R and K179R showed no difference. These findings indicate that the acetylation-SUMOylation balance at the RORβ hinge modulates transcription of a physiological relevant target gene, extending beyond effects observed with exogenous reporter constructs.

These findings have important implications for disease contexts in which RORβ functions. A regulatory circuit in which pharmacological modulation of SIRT1 shifts RORβ between stabilized and degraded states provides a framework for tuning receptor abundance in either direction, depending on tissue context and functional demand. Because the K176/K179 module resides within the hinge region rather than the ligand-binding pocket, it may offer a strategy to modulate RORβ activity without requiring a conventional ligand for this orphan receptor. This feature is particularly relevant to conditions such as retinal degeneration, osteoarthritis, RORβ-associated cancers, and the neurological disorders described above.

Several questions remain open. The structural basis of the RORβ-p300-SIRT1 ternary complex has not been resolved at high resolution, and the mechanism by which p300 displaces UBR5 remains unclear. In Addition, both K176R and K176Q exhibited elevated levels of non-SUMOylated RORβ even under PIAS1-driven conditions in which their SUMOylation was comparable to wild-type, raising the possibility that K176 contributes to receptor stability through mechanisms not fully captured by the SUMOylation circuit described here. Finally, the present work was conducted largely in transfected cell systems where the system can be defined; whether the K176-Ac/K179-SUMO axis operates with similar dynamics on endogenous RORβ in neurons, retinal cells, or osteoblasts, and whether it is altered in the disease contexts described above, remain important questions for future investigation. In summary, these findings identify a p300-SIRT1-PIAS1 axis, organized around the RORβ K176/K179 hinge module, as a determinant of RORβ stability and transcriptional output, and provide a framework for investigating whether dysregulation of this axis contributes to the disease contexts in which RORβ abundance is perturbed.

## Acknowledgements

The authors acknowledge the expert assistance of the Recombinant Protein Production and Characterization Core (RPPCC) at Baylor College of Medicine for expression of recombinant proteins in the baculovirus insect cell system. The RPPCC is supported by the NCI Cancer Center Support Grant (P30CA125123) of the Dan L. Duncan Comprehensive Cancer Center. Myc-p300/pCMVβ was a gift from Tso-Pang Yao (Addgene #30489).

## Author Contributions

**T.R.O.,** performed RIME, IP-MS, Western blots, and XL-MS experiments, analyzed data, wrote the paper. **D.S.,** performed mass spectrometric data acquisition and reporter assays. **N.I.M.,** performed qPCR, IP-MS, and reporter assays**. N.M.A.Y.,** performed IP-MS experiments and reporter assays. **V.D.,** contributed to cell culture and transfection experiments. **D.P.E.,** provided recombinant p300 and contributed to experimental design. **P.R.G.,** supervised the study, contributed to experimental design, analyzed the data, provided financial support and wrote the paper. **M.R.C.,** conceived and supervised the study, contributed to experimental design, analyzed the data, and wrote the paper.

## Declaration of generative AI and AI-assisted technologies in the manuscript preparation process

During the preparation of this work, the authors used Claude (Anthropic) and Co-Pilot (Microsoft) for editing purposes. The author(s) reviewed and edited the output as needed and take full responsibility for the content of the published article.

## Data Availability

These data have been deposited to the ProteomeXchange Consortium via the PRIDE^40^ partner repository with the dataset identifier PXD058187. The reviewer login details are as follows*:

Username:

Password:

*To be removed before final publication

## STAR Methods

### Key resources table

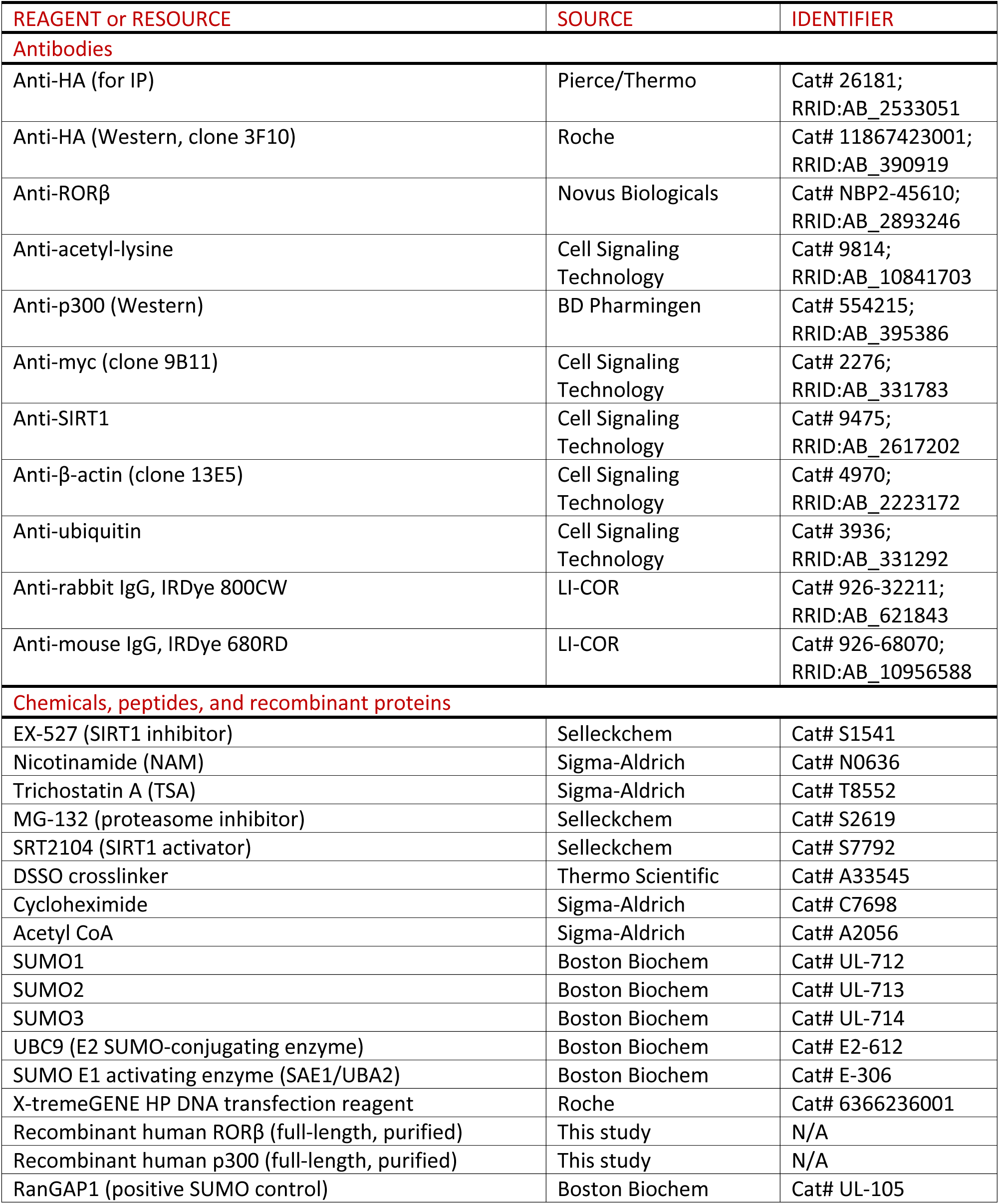
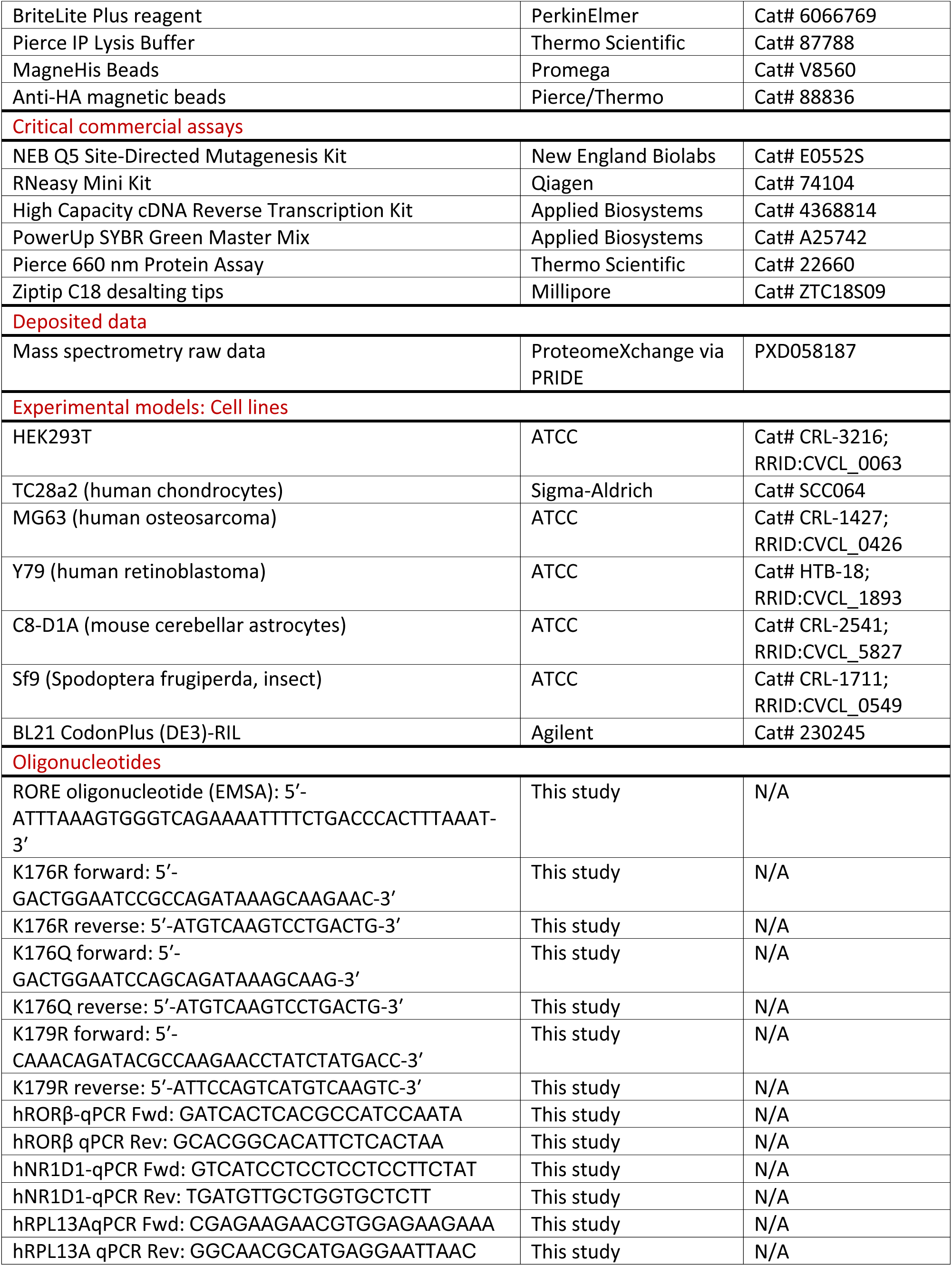
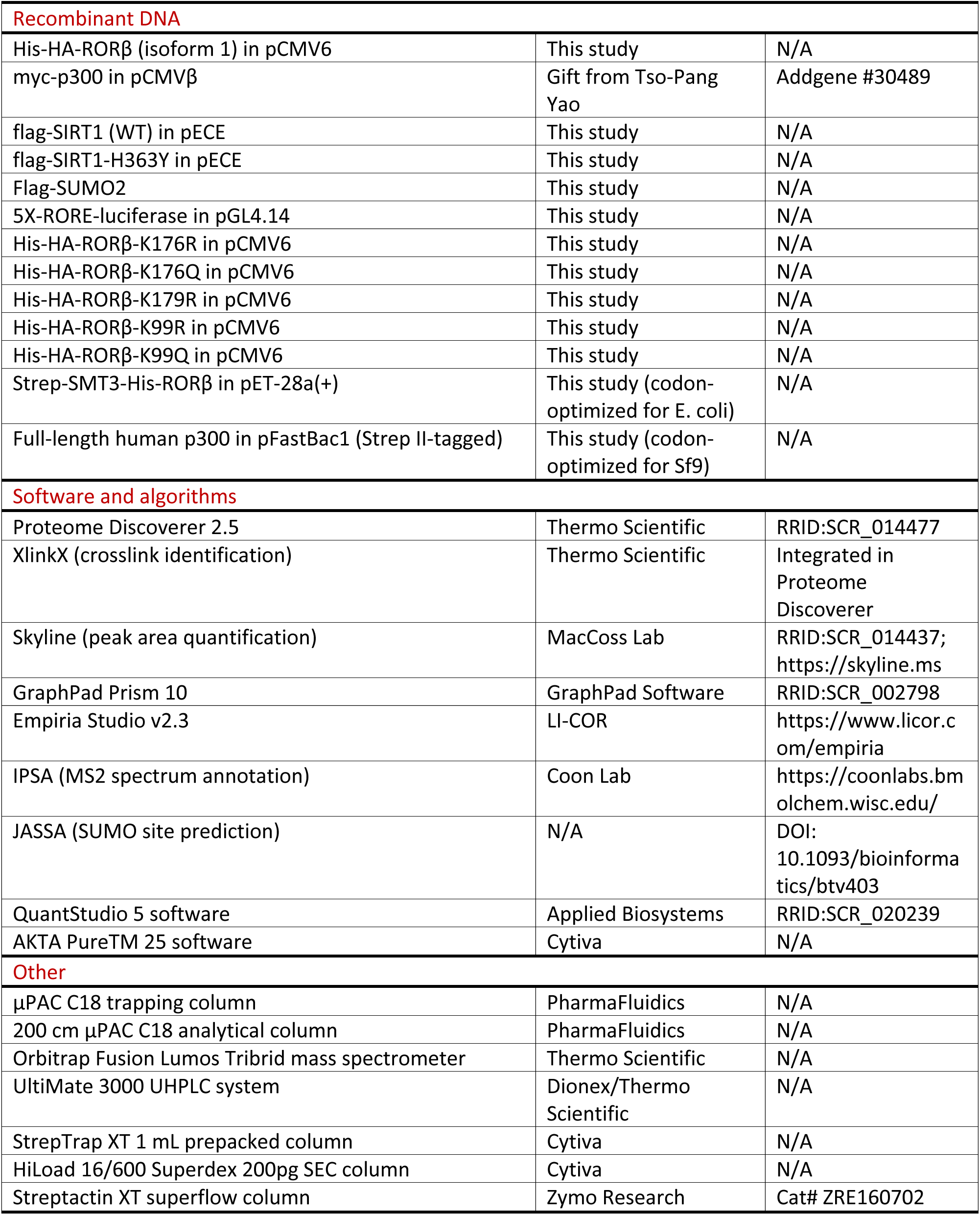

## Resource Availability

### Lead Contact

Further information and requests for resources and reagents should be directed to and will be fulfilled by the lead contacts: Patrick R. Griffin and Mi Ra Chang

## Materials Availability

Plasmids generated in this study will be deposited to Addgene upon publication. All unique reagents generated in this study are available from the lead contacts upon reasonable request.

## Data and Code Availability

All mass spectrometry raw data have been deposited in the ProteomeXchange Consortium via the PRIDE partner repository. This paper does not report original code. Any additional information required to reanalyze the data reported in this paper is available from the lead contacts upon reasonable request.

## Experimental Model Details

### Cell line and transfection

For rapid immunoprecipitation mass spectrometry of endogenous proteins (RIME) experiments, TC28a2 human chondrocytes (Sigma) were transfected with human His-HA-RORβ (or pCMV6 mock vector for control samples) using X-treme Gene HP DNA reagent (Roche) at 70 to 80% confluency at a ratio of 3:1 reagent to DNA. Cells were harvested 24 hours after transfection. RORβ overexpression (OE)-RIME was performed in TC28a2 cells. MG63 and Y79 cells were subjected to RIME experiments to identify endogenous RORβ- binding proteins. HEK 293T (ATCC) cells were used for Western blots, RORE-luciferase reporter assays, and mass spectrometry analysis of acetylation, SUMOylation and ubiquitination. HEK293T cells were transfected at a 1:2:3 ratio of His-HA-RORβ/pCMV6, myc-p300/pCMVβ, flag-SIRT1/pECE. Cells were transfected with X-tremeGENE HP reagent at 70 to 80% confluency at a ratio of 3:1 transfection reagent to DNA. Total transfected DNA was held constant by adding empty vector. myc- p300 /pCMVβ was a gift from Tso-Pang Yao (Addgene plasmid #30489). Mycoplasma tests were performed every 6 months. TC28a2, MG63, and HEK 293T cells were maintained in Dulbecco’s modified Eagle’s medium (DMEM) supplemented with 10% fetal bovine serum (FBS) with antibiotics (penicillin/streptomycin; Invitrogen). Y79 cells were maintained in RPMI-1640 with 10% FBS and antibiotics.

### Luciferase reporter assay

Twenty-four hours post-transfection, cells were replated into a 384-well plate at a seeding density of 5,000 cells/well in 20 µL of complete medium. HEK 293T cells were incubated for 4-5 h at 37 °C to allow adherence, after which cells were treated with either DMSO or compounds. Following this, the Dual-Glo® Luciferase Assay (Promega) was performed according to the manufacturer’s instructions. In brief, plates were equilibrated to room temperature, and an equal volume of Dual-Glo® Luciferase Reagent was added to each well. After 10 min of incubation with shaking (3 min at 600 rpm), firefly luciferase activity was measured. An equal volume of Dual-Glo® Stop & Glo® Reagent was then added, and after a further 10 min of incubation at room temperature, Renilla luciferase activity was measured. Luminescence was measured on a Synergy Neo2 microplate reader (BioTek) with an integration time of 0.5 s and automatic gain adjustment. Results were analyzed using GraphPad Prism.

### RIME (Rapid Immunoprecipitation Mass Spectrometry of Endogenous Proteins)

RIME experiment was performed in cells transfected with HA-tagged RORβ using an adapted protocol for formaldehyde crosslinking of chromatin complexes followed by immunoprecipitation^35^. Approximately 1.5 x 10^7^ cells were used per bioreplicate (n=3). 24 hours after transfection, cells were washed with PBS and media was replaced with freshly prepared room temperature PBS containing 1% formaldehyde (Thermo Scientific, 28908). After 8 minutes at room temperature, the reaction was quenched by adding glycine (1.0 M, pH 7.5) to a final concentration of 100 mM. Cells were washed twice with cold PBS, harvested, and treated with lysis buffer to remove cytoplasmic proteins (50 mM HEPES-KOH, (pH 7.5), 140 mM NaCl, 1 mM EDTA, 10% (vol/vol) glycerol, 0.5% (vol/vol) NP-40/Igepal CA-630 and 0.25% (vol/vol) Triton X-100 with HALT protease inhibitor (Thermo)), nuclei were pelleted by centrifugation and treated with lysis buffer 2 (10 mM Tris-HCl (pH 8.0), 200 mM NaCl, 1 mM EDTA and 0.5 mM EGTA, with HALT protease inhibitor). After centrifugation, pellet was resuspended in lysis buffer 3 (10 mM Tris-HCl (pH 8.0), 100 mM NaCl, 1 mM EDTA, 0.5 mM EGTA, 0.1% (wt/vol) sodium deoxycholate and 0.5% (vol/vol) N-laurylsarcosine) and DNA was fragmented with sonication (Corvaris). Triton X-100 was added to 1% and samples were centrifuged (20,000 x *g*, 10 min, 4 °C). Supernatant was mixed with anti-HA magnetic beads (Pierce), 33 µg HA peptide in Pierce IP/lysis buffer was added to negative control samples (with corresponding vehicle added to experimental samples) and incubated at 4 °C overnight with rotation. Beads were washed nine times (50 mM HEPES (pH 7.6), 1 mM EDTA, 0.7% (wt/vol) sodium deoxycholate, 1% (vol/vol) NP-40 and 0.5M LiCl) followed by two washes with 100 mM ammonium bicarbonate. Beads were digested with trypsin overnight at 37 °C (0.5 µg trypsin, 100 mM ammonium bicarbonate). Following trypsin digestion, peptides were desalted with ziptip (Millipore, ZTC18S09) following the manufacturer’s protocol.

### Expression and purification of recombinant RORβ

Strep-SMT3-His-RORβ was codon optimized for *E. Coli* expression and inserted into a pET-28a(+) plasmid (GenScript). After transformation (Agilent, BL21 CodonPlus (DE3)-RIL, 230245) cells were grown on agar supplemented with kanamycin and single colonies were used to prepare overnight culture in terrific broth with kanamycin. Overnight culture (0.5 mL) was added to 250 mL terrific broth media (with kanamycin) containing 30 µM ZnCl_2_, grown at 37 °C to an OD_600_ of 0.6-0.8, and the temperature was reduced to 16 °C. IPTG was added to 250 µM and the media was shaken for an additional 15 h. Cells were harvested (4,000 x *g*, 20 min, 4 °C) and lysed (50 mM HEPES, pH 8.0, 500 mM NaCl, 10% glycerol, 1 mM DTT, 1 mM MgCl_2_, with 0.25 mg/mL lysozyme, 400 units benzonase, with HALT protease inhibitor and 1 mM PMSF, 30 mL total volume, 30 minutes on ice) followed by 2 minutes sonication at 70% amplitude. Sample was clarified (20,000 x *g*, 15 minutes, 4 °C) and flowed over 1.1 mL volume streptactin column (Streptactin XT superflow, Zymo Research, ZRE160702). Flow through was collected and added to the column a second time, then the column was washed with 5 column volumes of wash buffer (50 mM HEPES, pH 8.0, 500 mM NaCl, 10% glycerol). SUMO protease (Ulp1) was added (1.0 µg in 1.0 mL 50 mM HEPES, pH 8.0, 500 mM NaCl, 10% glycerol, 5 mM DTT) and incubated 1 h at RT. Sample was eluted in 3.0 mL (50 mM HEPES, pH 8.0, 500 mM NaCl, 10% glycerol, 5 mM DTT) and stored in 50% glycerol at -20 °C.

### Expression and purification of recombinant p300

Human p300 as full-length open reading frame DNA was synthesized with optimal codon usage for insect cells with an in-frame Strep II tag (WSHPQFEK/G) and a glycine spacer at the amino-terminus and inserted into pFastBac1 transfer vector (Epoch Life Sciences, Houston, TX). Recombinant bacmid was generated and expanded in Spodoptera frugiperda (Sf9) cell cultures and viral titer was determined by plaque assays as previously described.^38, 39^ Multiple 500 mL cultures of SF9 cells were infected with recombinant virus at an MOI of 2.0 and incubated for 48 hour at 27°C in oxygenated spinner vessels. Cells were collected and centrifuged 1500x g for 10 min and pellets were washed once PBS by resuspension and centrifugation. Cell pellets were flash frozen and stored at -80°C. Cells (from 2x 500ml cultures) were resuspended in 50 mL of lysis buffer (50 mM Na_2_HPO_4_, pH 8.0, 500 mM NaCl, 5% Glycerol, 1 M Urea, 1 mM EDTA, 1 mM 2ME), supplemented with protease inhibitor tablets (leupetin, aprotinin, bacitracin, and PMSF) and submitted to Teflon-glass homogenization for 8 strokes at 1.5 speed in the cold room (4°C). The homogenate was treated with 1.2 units/mL of benzonase nuclease or 1 hour at 4°C then passed 3 times through an 18G needle followed by a 25G needle. The lysate was centrifuged twice for 1 hour at 50,00 xg and the resulting supernatant was filtered using 0.2 μm disposable filters. Purification was performed on AKTA Pure^TM^ 25 at 4 °C while monitoring conductivity and UV. A 1 mL StrepTrap XT prepacked Hi-Trap chromatography column (Cytiva) was equilibrated with lysis buffer for 10 column volumes (CV), and after loading the cleared cell lysate, the column was washed for 30 CV with equilibration buffer and then eluted with equilibration buffer containing 40 mM biotin. Eluted fractions (1 mL) were collected and analyzed by Coomassie Blue-stained 4-15% gradient SDS-PAGE gel. Fractions containing p300 were pooled and concentrated to 1-2 mL by an Amicon ultracentrifugal device with a 10,000 MW cutoff (A280<4.0). Protein was further purified by a preparative size exclusion chromatography (SEC) column (HiLoad 16/600 Superdex 200pg), degassed and equilibrated in SEC buffer (20 mM Hepes, pH 7.5, 200 mM NaCl, 5% Glycerol, 1 M Urea and 1 mM TCEP). Elution fractions (1 mL) were collected and analyzed by Coomassie Blue-stained 4-15% gradient SDS-PAGE gels and peak fractions were concentrated as above. Concentration of p300 was by determined by Nanodrop absorbance at 280 nm, calculation of extinction coefficient, and by comparison of purified bands on Coomassie Blue-stained 4-15% SDS-PAGE gel with a standard curve of know amounts of unstained protein markers. Aliquots (50-100 μL) of p300 were snap frozen in liquid nitrogen and stored at -80°C for up to 3-4 months and samples were used only once after thawing.

### Co-immunoprecipitation experiments

For RORβ-p300 Co-IP, TC28a2 cells were transiently transfected with HA-RORβ and myc-p300 or base vector with xtremegene HP at 4:1 ratio of p300 to RORβ. After 24 h the cells were treated with 400 nM TSA for 20 h. Cells were harvested and lysed for 30 minutes on ice after resuspension in lysis buffer (250 mM NaCl, 0.1% NP-40, 20 mM NaH2PO4, pH 7.5, 5 mM EDTA, 30 mM sodium pyrophosphate, 10 mM NaF, 5 mM nicotinamide, 400 nM TSA, with HALT protease inhibitor (Thermo Fisher Scientific)). Protein concentration was assessed by BCA and protein concentrations were normalized across conditions. HA-RORβ was immunoprecipitated with anti-HA beads (Pierce) and washed twice with 300 uL of wash buffer (10 mM HEPES, 100 mM NaCl, 0.5 mM EDTA, 0.05% tween-20). Beads were eluted with 50 µL of nonreducing SDS-PAGE loading buffer by heating at 95 °C for 5 minutes, then reduced an additional 2 minutes at 95 °C with 25 mM DTT. Gels were loaded with 10 µg lysate and 20 µL of eluate and transferred to nitrocellulose membrane. For RORβ-SIRT1, HEK 293T cells were transfected 1:1 with RORβ and Flag-SIRT1 with PEI reagent. After 48 hours, the transfected cells were harvested and treated as above, except three washed were done, and for negative control samples 75 µg of HA peptide were added to each 900 µL negative control sample.

### CHX protein stability

HEK 293T cells were transfected and after 42 h, cycloheximide (CHX) was added at 10 µM at intervals of 4, 2, 1, and 0 h (0 h treated with DMSO only). Cells were washed with PBS, harvested and lysed (250 mM NaCl, 0.1% NP-40, 20 mM NaH2PO4, pH 7.5, 5 mM EDTA, 30 mM sodium pyrophosphate, 10 mM NaF with HALT protease inhibitor (Thermo Fisher Scientific)), and centrifuged (16,000 x *g*, 10 min, 4 °C). Protein concentration was measured with Pierce 660 assay (Thermo Scientific), and 10 µg protein was loaded per well for SDS-PAGE (n=2). Following transfer, membranes were immunoblotted (RORβ (Novus, NBP2-45610), β-actin (Cell Signaling, 13E5) 1000:1 ratio, overnight, 4 °C). Membranes were washed (3 x 5 min, TBST), treated with secondary antibodies (Li-COR, anti-rabbit 926-32211, anti-mouse 926-68070, 15,000:1, 1 h, RT) and imaged (Li-Cor). Fluorescence signal was quantified with Empiria Studio software (Version 2.3.0.154) and data was analyzed in GraphPad Prism (Version 6.01).

### *In cellulo* RORβ acetylation, ubiquitination and SUMOylation analysis

RORβ acetylation was analyzed by adapting a prior method ^42^ HEK 293T cells were transfected and after 30 min, trichostatin A (TSA) was added to a concentration of 400 nM. After 24 h, cells were washed 1 x with ice cold PBS and harvested (approximately 1.5 x 10^7^ cells). Cells were lysed (250 mM NaCl, 0.1% NP-40, 20 mM NaH2PO4, pH 7.5, 5 mM EDTA, 30 mM sodium pyrophosphate, 10 mM NaF with HALT protease inhibitor (Thermo Fisher Scientific), 400 nM TSA, 5 mM NAM) for 20 min on ice. Sample was clarified and incubated with 100 µL anti-HA magnetic beads (Pierce, 45 min, RT), washed (5 x, 1 mL IP/lysis buffer (Pierce) supplemented with additional 100 mM NaCl). Sample was eluted (95 °C, 5 min, 25 µL non-reducing lane marker (Thermo Scientific) and loaded onto SDS-PAGE gel. Bands for RORβ were excised from the gel, washed (200 µL 1:1 acetonitrile/50 mM ammonium bicarbonate, 20 min, 1000 rpm, followed by 50 mM ammonium bicarbonate, 20 min, 1000 rpm). Band was dehydrated (100 µL acetonitrile, 5 min speedvac), then reduced (200 µL, 25 mM DTT, 30 min, 56 °C), alkylated (200 µL, 55 mM iodoacetamide, in dark, 30 min), washed twice (400 µL water), dehydrated, and digested with trypsin (0.1 µg, 200 uL 50 mM ammonium bicarbonate, 18h). Peptides were extracted twice (1:1 acetonitrile / 0.1% TFA, 20 min), dried, and desalted with ziptips (Millipore, ZTC18S09).

To search for ubiquitin modifications a similar immunoprecipitation strategy was employed. Proteins were digested on-bead with a combination of trypsin and chymotrypsin. Four hours prior to harvesting cells, MG-132 (or DMSO) were added to cell media to a concentration of 20 µM. A low concentration of iodoacetamide (1 mM) was included during cell lysis and immunoprecipitation at 4 °C to inhibit deubiquitinases, but samples were not reduced and alkylated to avoid high concentrations of iodoacetamide which could acylate lysine residues ^43^. For MS detection of the modification sites, data was analyzed in Proteome Discoverer.

### Liquid chromatography and tandem mass spectrometry

Samples were injected (duplicate or triplicate injections) onto an UltiMate 3000 UHP liquid chromatography system (Dionex, ThermoFisher). Peptides were trapped using a μPAC C18 trapping column (PharmaFluidics) using a load pump operating at 20 μL/min. Peptides were separated on a 200 cm μPAC C18 column (PharmaFluidics) using a linear gradient (1% Solvent B for 4 min, 1–30% Solvent B from 4 to 70 min, 30–55% Solvent B from 70 to 90 min, 55–97% Solvent B from 90 to 112 min, and isocratic at 97% Solvent B from 112 to 120 min) at a flow rate of 800 nL/min. Gradient Solvent A contained 0.1% formic acid, and Solvent B contained 80% acetonitrile and 0.1% formic acid. Liquid chromatography eluate was interfaced to an Orbitrap Fusion Lumos Tribrid mass spectrometer (ThermoFisher) with a Nanospray Flex ion source (ThermoFisher). The source voltage was set to 2.5 kV, and the S-Lens RF level was set to 30%. Full scans were recorded from m/z 150 to 1500 at a resolution of 60,000 in the Orbitrap mass analyzer. The AGC target value was set to 4 × 10^5^, and the maximum injection time was set to 50 ms in the Orbitrap. For PTM analysis, MS2 scans were recorded at a resolution of 15,000 in the Orbitrap mass analyzer. The AGC target was set to 5 × 10^4^, with a maximum injection time of 180 ms, and an isolation width of 1.6 m/z. For RIME analyses, MS2 was performed in the iontrap, with an isolation width of 1.2 m/z using a turbo scan rate and dynamic maximum inject time. FAIMS compensation voltages were set on –45 and –60 V. Cross-links were identified using a previously described MS2-MS3 method with slight modifications ^44^. Full scans were recorded from m/z 150 to 1500 at a resolution of 60 000 in the Orbitrap mass analyzer. The AGC target value was set to 4 × 10^5^, and the maximum injection time was set to 50 ms in the Orbitrap. MS2 scans were recorded at a resolution of 30 000 in the Orbitrap mass analyzer. Only precursors with a charge state between 3 and 8 were selected for MS2 scans. The AGC target was set to 5 × 10^4^, a maximum injection time of 150 ms, and an isolation width of 1.6 m/z. The CID fragmentation energy was set to 25%. The two most abundant reporter doublets from the MS2 scans with a charge state of 2–6, a 31.9721 Da mass difference, and a mass tolerance of ±10 ppm were selected for MS3. The MS3 scans were recorded in the ion trap in rapid mode using HCD fragmentation with 35% collision energy.

### Cross-linking Mass Spectrometry (XL-MS)

Crosslinking of His-RORβ with p300 was performed with 5 µM of each protein with DSSO crosslinker (2.0 mM DSSO, 1 h, RT. buffer: HEPES 50 mM, pH 7.5, 150 mM NaCl). Crosslinking reactions were performed in triplicate for both DSSO and DMSO controls. Putative crosslinks were identified in Proteome Discoverer with the XlinkX algorithm workflow.

### RORβ *in vitro* acetylation by p300 and EMSA

Full length recombinant human RORβ was incubated with full length recombinant human p300, with 1 mM acetyl CoA (Sigma) at a 20:1 w/w ratio of RORβ to p300 (HEPES 50 mM, 10 mM sodium butyrate, 7.5% glycerol, 1 mM DTT, HALT protease inhibitor, pH 8.0; 1 h at 30 °C). Reactions were carried out with RORβ at concentrations of 4.0 µM and 2.0 µM. For DNA binding EMSA assay, the above acetylation reactions were incubated with DNA (0.25 µM) in ratios of 8:1 and 4:1 RORβ to DNA (sequence 5’-ATTTAAAGTGGGTCAGAAAATTTTCTGACCCACTTTAAAT-3’) respectively (HEPES 25 mM, 5 mM sodium butyrate, 3.75% glycerol, 0.5 mM DTT, 200 mM NaCl, 10 mM MgCl2, HALT protease inhibitor, pH 8.0; 20 min, RT) and assayed on an 8% nondenaturing polyacrylamide gel (200 V, 40 min, 4 °C).

### Mutant RORβ Generation

RORβ mutants K176R, K176Q, and K179R were generated from human His-HA-RORβ in pCMV6 base vector with using the primers K176R forward (5’-GACTGGAATCCGCCAGATAAAGCAAGAAC-3’) K176R reverse (5’-ATGTCAAGTCCTGACTG-3’), K176Q forward (5’-GACTGGAATCCAGCAGATAAAGCAAG-3’) and K176Q reverse (5’-ATGTCAAGTCCTGACTG-3’), K179R forward (5’-CAAACAGATACGCCAAGAACCTATCTATGACC-3’) and K179R reverse (5’- ATTCCAGTCATGTCAAGTC-3’) using NEB Q5 site-directed mutagenesis kit (#E0552). The K99R mutant in the same mammalian expression vector was prepared using the In-Fusion cloning method (Takara) with primers: K99R forward (5’- GGTGCAGAGACACCAGCAGCGGCTGCAG-3’) and K99R reverse (5’- TGGTGTCTCTGCACCTCAGCATACAGGCTG-3’). Mutants K176R and K179R for recombinant RORβ in pET-28a(+) vector were also prepared using the In-Fusion cloning method. The primers used were K176R forward (5’-CGGCATCCGTCAAATTAAACAGGAGCCGATTTACG-3’), K176R reverse (5’-ATTTGACGGATGCCGGTCATATCCAGAC-3’), K179R forward (5’-GCAAATTCGTCAGGAGCCGATTTACGATCTGACC-3’) and K179R reverse (5’-TCCTGACGAATTTGCTTGATGCCGGTCATATCC-3’).

### RT-PCR

MG-63 cells were transfected with 2 ug of wild-type (WT), base vector (NC) or mutant RORB plasmid using X-tremeGENE HP DNA Transfection Reagent (Roche) at a 3:1 reagent:DNA ratio. After 48 hours, cells were collected and harvested for total RNA using the RNeasy Mini Kit (Qiagen). RNA was reverse transcribed using the High Capacity cDNA Reverse Transcription Kit (Applied Biosystems/Thermo Fisher Scientific). Quantitative PCR was performed with a QuantStudio 5 Real-Time PCR System (Applied Biosystems/Thermo Fisher Scientific) using PowerUp SYBR Green Master Mix (Applied Biosystems/Thermo Fisher Scientific).

### Data deposition

All MS data has been submitted to the ProteomeXchange Consortium via the PRIDE partner repository with the dataset identifier, PXD058187.

