## Supplemental Figures for "Acetylation-Primed SUMOylation Drives RORβ Turnover via a p300-SIRT1 Regulatory Axis"

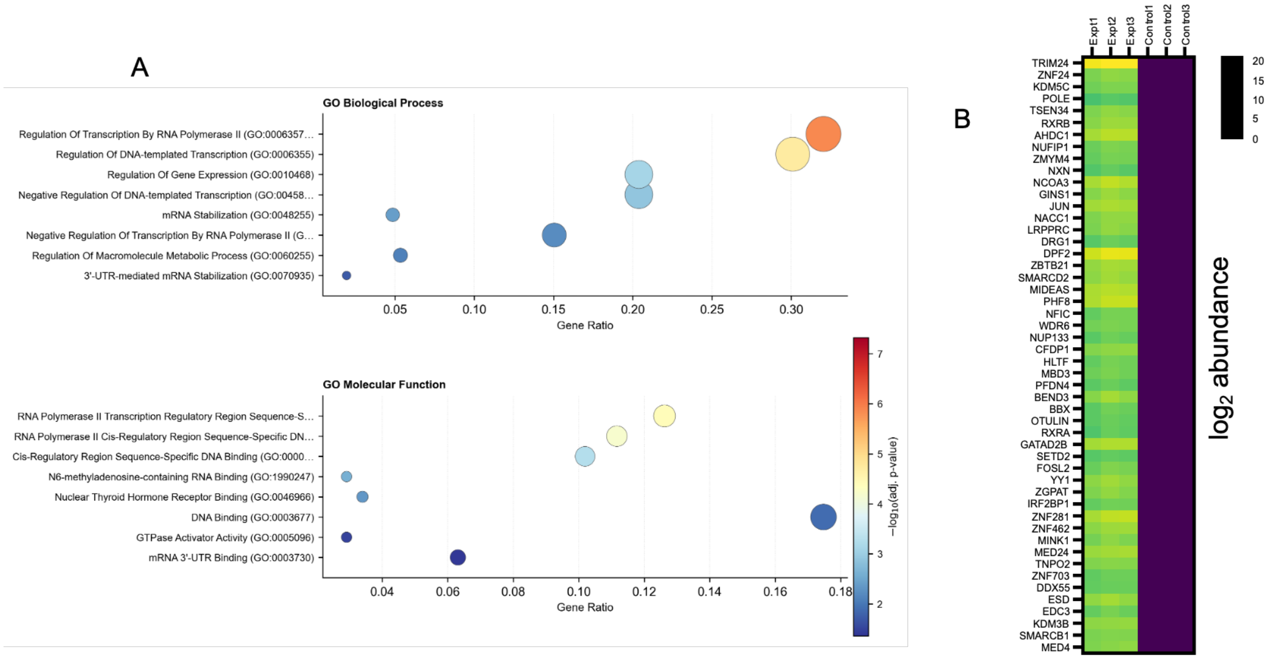


**Figure S1**. A. GO analysis of 206 RORβ interactors found in common for two full repeated RIME experiments. The list of 206 proteins was taken from proteins found in both datasets with log2FC ≥ 1.5 and p-value < 0.05. GO analyses were performed with David Bioinformatics tools (DOI: 10.1093/bioinformatics/bth456). B. Heatmap of top 50 highest confidence RORβ interactors sorted by p-value (with lowest p-values ranked highest). This corresponds to the experimental dataset shown in **Fig. 1A**.

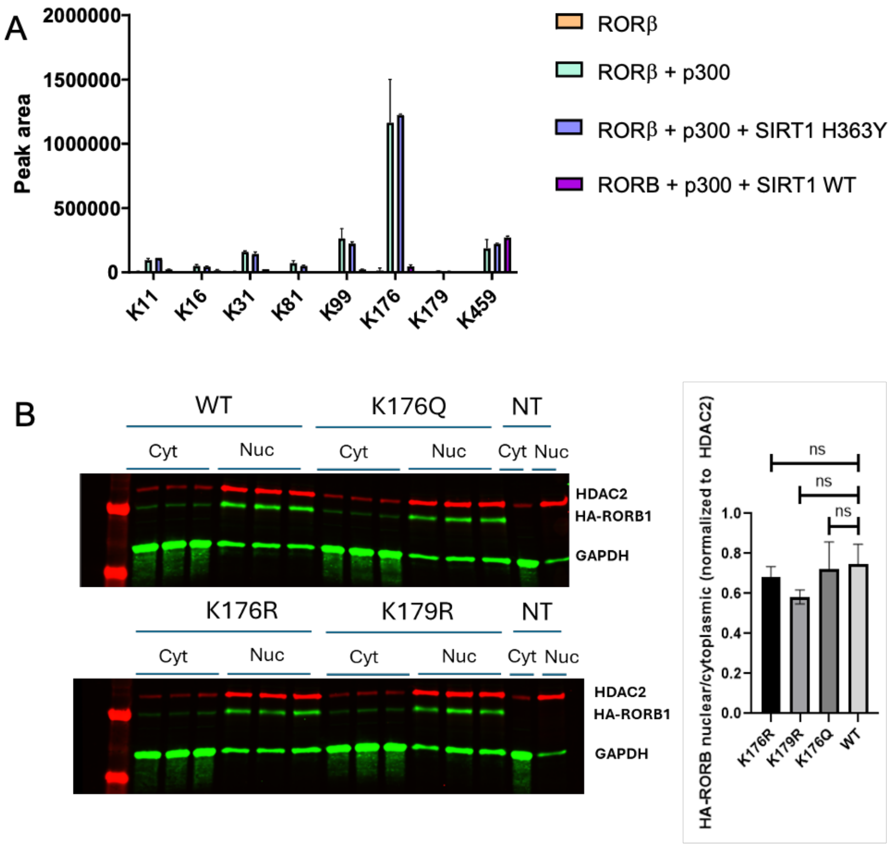


**Figure S2**. A. RORB acetylation sites detected with trypsin/chymotrypsin digest and including conditions with the catalytically inactive SIRT1 H363Y mutant. Following IP-MS of RORβ and after washing the beads the protease digest was performed on-bead. B. Subcellular fractionation of HEK293T transiently transfected cells shows no significant differences in nuclear/cytoplasmic partitioning. After nuclear/cytoplasmic partitioning samples were immunoblotted for HA and for the nuclear marker HDAC2 and cytoplasmic marker GAPDH.

A

RORβ SUMO motif analysis

| Results for putatifs SUMO site | | | | | | | | |
|---|---|---|---|---|---|---|---|---|
| Position K | Sequence | Best PS | Consensus direct | | | Consensus Inverted | | |
| | | | Type | PSd | DB Hit | Type | PSi | DB Hit |
| K16 | EVIPCKICGDKSSGIHYGVIT | None | None | None | 1 | None | None | |
| K81 | LALGMSRDAVKFGRMSKKQRD | None | None | None | 1 | None | None | |
| K151 | YANGHVIDLPKSEGYYNVDSG | High | Consensus | High | 1 | None | None | |
| K176 | DQSGLDMTGIKQIKQEPIYDL | None | None | None | | None | None | 4 |
| K179 | GLDMTGIKQIKQEPIYDLTSV | High | SC-SUMO | High | 57 | None | None | 7 |
| K248 | AWQTHTYEEIKAYQSKSREAL | Low | None | None | | Consensus inv | Low | |
| K278 | HAIQYVVEFAKRITGFMELCQ | None | None | None | | None | None | 2 |
| K296 | LCQNDQILLLKSGCLEVVLVR | None | None | None | | None | None | 4 |
| K323 | PLNNTVLFEGKYGGMQMFKAL | None | None | None | | Weak consensus inv | None | |
| K380 | PDRAWLIEPRKVQKLQEKIYF | None | None | None | 1 | None | None | |
| K408 | KNHLDDETLAKLIAKIPTITA | None | None | None | | None | None | 1 |
| K426 | ITAVCNLHGEKLQVFKQSHPE | None | None | None | | None | None | 1 |

B

human RORβ NCBI BLASTP to 1000 animal RORβ sequences

| Site | Mismatches (%) |
|---|---|
| G174 | 19.3 |
| I175 | 48.2 |
| K176 | 11.7 |
| Q177 | 76.4 |
| **I178** | 0.5 |
| **K179** | 0.0 |
| **Q180** | 0.0 |
| **E181** | 0.0 |
| P182 | 18.1 |
| I183 | 27.6 |

C

In vitro RORB SUMOylation: IB for SUMO1

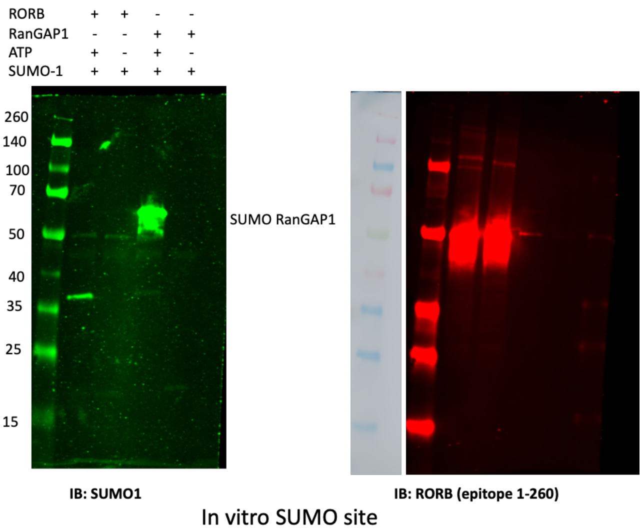


**Figure S3**. A. SUMO motif analysis of RORβ amino acid sequence was performed with Jassa webserver (DOI: 10.1093/bioinformatics/btv403). and shows a SUMO motif at K179 with 57 matches to known SUMO sites. B. Conservation analysis of select residues in the RORβ hinge was performed by a multiple sequence alignment of 1000 animal RORβ sequences using NCBI BLASTP. Conservation was assessed by the rate of mismatches at the residues shown. C. In vitro SUMOylation assay for RORβ conjugation to SUMO1. Recombinant RORβ was incubated with E1 and E2 ligases and SUMO1, +/- ATP required for SUMOylation. RanGAP1 served as a positive control. Samples of the reactions were immunoblotted for SUMO1 and RORβ.

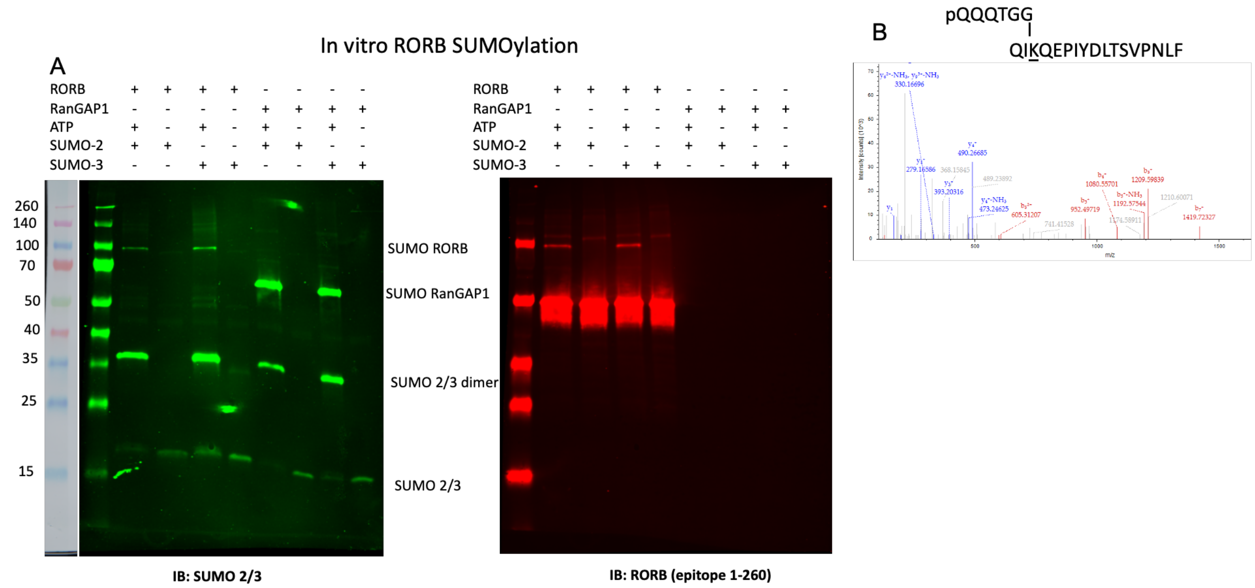


**Figure S4**. A. In vitro SUMOylation assay for RORβ conjugation to SUMO2 and SUMO3. Recombinant RORβ was incubated with E1 and E2 ligases and SUMO2 or SUMO3, +/- ATP required for SUMOylation. RanGAP1 served as a positive control. Blotting was done for SUMO2/3 or for RORβ. B. Mass spectrometry MS2 assignment of K179-containing peptide with QQQTGG SUMO2 remnant. Spectra taken from reaction with SUMO2 after excising gel band at 100 kDa. The “p” on the N-terminal Q of the QQQTGG sequence represent pyro- modification of this residue, a common artifact in mass spectrometry of peptides with an N-terminal Q.
